# Geometry-Conditioned Hyperalignment for Generalizable EEG Decoding Across Datasets and Paradigms

**DOI:** 10.64898/2026.11.08.765221

**Authors:** Huu Hoang, Shanglin Li, Okito Yamashita, Motoaki Kawanabe, Mitsuo Kawato

## Abstract

Electroencephalography (EEG) brain-computer interfaces (BCIs) supports promising applications, but daily use remains limited by poor cross-subject and cross-dataset generalisation. That is because EEG recordings made with different electrode montages do not even share an observation space, and each subject’s leadfield projects the same neural sources onto the scalp differently. We present **hTSMNet**, which combines a novel geometry-conditioned hyperalignment frontend with a Riemannian symmetric positive-definite (SPD) manifold backbone that reduces the residual cross-subject distribution shift. From electrode coordinates alone, the frontend builds a linear map from any montage onto a fixed set of scalp *anchors*, placing every recording in one shared space. Because this map has no subjectspecific parameters, an unseen subject, dataset, or montage is aligned from its geometry alone, so transfer is zero-shot by construction. On a leave-one-dataset-out left-/right-hand motor-imagery (MI) benchmark across three datasets (172 subjects) and eight comparators, hTSMNet matches the strongest comparator at strict zero-shot, and attains the highest accuracy on every dataset once a few calibration trials for adjusting only final-layer classifier are available. We show that it orients the binary decision the same way in every dataset, so that a single linear rule learned on the source data is already correctly oriented on a held-out one. The same *frozen* MI-pretrained model also transfers across *paradigms*: with only a re-fit final-layer classifier, hTSMNet decodes mental workload and steady-state visual evoked potentials (SSVEP) well above chance, yet remains at chance on stimulus-locked event-related potentials (ERP). This dissociation follows from the architecture: covariance pooling reads oscillatory band power and is largely insensitive to the phase-locked potentials that ERP tasks depend on. What the representation generalizes is a *signal family* (band-limited oscillatory activity), not the named task. Geometry-conditioned hyperalignment thus yields an inspectable representation, which is generalizable across datasets and, within the oscillatory regime, across paradigms.

## 1 Introduction

Electroencephalography (EEG) brain-computer interfaces (BCIs) support promising applications in communication, rehabilitation, cognitive monitoring, and passive-state sensing [1, 18, 31], but routine use is still limited by poor generalisation across subjects, recording sites, and datasets [20]. A central obstacle to generalizable EEG decoders is that the measurement space is not shared across subjects, laboratories, or datasets. Even when two recordings probe the same neural process, they may differ in electrode count, cap layout, reference scheme, and sampling convention. More fundamentally, each subject has a distinct forward operator: the same latent source activity produces different scalp patterns because of differences in head geometry, electrode placement, and tissue conductivity. Cross-dataset transfer is therefore not a problem of statistical distribution shift alone. It is a problem of *incompatible observation spaces*. The issue is acute in motor-imagery (MI) EEG, where public benchmarks span small 10–20 montages to dense 64-channel arrays [27, 28], but the same fundamental difficulty applies to MEG, intracranial local field potentials, and other variable-coverage electrophysiological recordings.

Several established lines of work address this heterogeneity, but each falls short of a general, geometry-based solution. They divide into three broad strategies. One practical solution forces every recording into a common input shape—selecting a shared electrode subset or interpolating each montage onto a fixed template—so that a conventional fixed-input, subject-invariant network can be trained across recordings [2]. This makes the data compatible with conventional networks, but the alignment is prescribed before learning and may not match the task-relevant neural geometry. A second line of work, domain adaptation by Euclidean alignment (EA) [3] or Riemannian Procrustes analysis (RPA) [4], targets the *statistical distribution shift* between domains by re-centring or aligning covariance statistics. This is effective when target data are available, but it is a different problem from montage incompatibility: these methods treat individual subjects as domains within a common channel space, draw on source and/or target data from that space, and are not intended to map heterogeneous montages onto one another. A third strategy trains variable-montage encoders that ingest channel tokens tagged with sensor position, as in recent EEG foundation models such as BIOT [8], LUNA [9], and LaBraM [10]. Earlier self-supervised models such as BENDR [7] scaled EEG pretraining but on a fixed montage. These models are powerful and scalable, but the mapping from sensors to a shared neural space is implicit in a large attention model rather than represented as an explicit, inspectable geometric operator.

Riemannian symmetric positive-definite (SPD)-manifold methods provide a complementary strength [5, 6]. Because covariance matrices of band-limited neural time series lie on the SPD manifold, tangent-space and SPD-network methods exploit the spatial covariance structure that is highly informative for MI decoding [11, 12]. Our previous work TSMNet [13] integrates temporal-spatial filtering, covariance pooling, SPD batch normalisation, and tangent-space classification, and partially reduces inter-subject variability by normalising covariance statistics on the manifold. Two gaps remain for cross-dataset use, however. First, TSMNet assumes a fixed channel count, so it cannot ingest heterogeneous montages directly. Second, its statistical normalisation does not explicitly account for the forward-model (leadfield matrix) differences that cause shared neural activity to manifest differently across montages. Closing both gaps calls for a module placed in front of the backbone: a montage-agnostic, geometry-aware frontend that re-expresses every recording in a single shared coordinate frame the covariance-aware decoder can consume, and that conditions this re-expression on electrode geometry rather than on covariance statistics alone.

We propose **hTSMNet** (hyperalignment TSMNet), which realises this frontend by defining a fixed set of *D anchor* locations on the scalp, into which each native montage is resampled. For each recording *s* with *N_s_* sensors, a geometry-conditioned frontend generates an alignment matrix *A_s_ ∈* ℝ*^N_s_×D^* that maps the native montage into this shared *D*-dimensional anchor space. The aligned signal 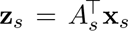 then feeds the TSMNet backbone, which always sees a fixed-dimensional input. The alignment *A_s_*is generated from the relative geometry between the recording’s electrodes and the anchors: a radial-basis-function (RBF) interpolation kernel refined by a shared, position-conditioned network and retracted onto a near-orthonormal projector. This gives the model a strong inductive bias— sensors should contribute strongly to nearby anchors—while leaving the precise mapping to be learned end-to-end from the task. Crucially, the frontend has *no subject-specific parameters*: the alignment of any new recording is determined by its sensor coordinates alone, so hTSMNet can transfer zero-shot to unseen subjects, datasets, and montages without target-domain calibration. We use *hyperalignment* in the sense of mapping heterogeneous recordings into one shared representational space. Unlike functional hyperalignment of fMRI responses [15], the map here is conditioned on electrode geometry rather than estimated from a shared stimulus-driven response, and so requires no target data.

Beyond the frontend architecture itself, we make three empirical contributions:

**1. Cross-dataset transfer under heterogeneous montages.** On a leave-one-dataset-out (LODO) MI left-/right-hand discrimination task across three datasets from MOABB (Mother of All BCI Benchmarks [20]), hTSMNet is compared with eight comparators spanning classical Riemannian pipelines, specialist deep networks, and pretrained foundation models (Section 2.3). hTSMNet matches the strongest comparators at strict zero-shot. Once a few final-layer classifier-only calibration trials are available, it obtains the highest few-shot accuracy on every dataset, with a per-subject advantage that is statistically significant against all eight comparators. This suggests that the SPD manifold representations of hTSMNet are generalizable across datasets because the frontend, spatiotemporal convolutions, covariance and tangent space mapping are not altered in the final-layer classifier calibration.
**2. A stable tangent-space representation.** We link this advantage to a geometric property of the learned representation. Rather than merely intermixing datasets in feature space, hTSMNet learns a left-/right-hand class direction shared across datasets, attaining the highest cross-dataset class-direction consistency of any method, so that a single linear decision rule is already oriented correctly on a held-out dataset (Section 2.4).
**3. Cross-paradigm transfer within the oscillatory regime.** The same shared geometry generalises beyond the training paradigm: a *frozen* MI-pretrained hTSMNet decodes mental workload (84%) and steady-state visual evoked potentials (SSVEP, 68–72%) well above chance from a re-fit linear classifier alone, while remaining at chance on stimulus-locked event-related potentials (ERP) (Sections 2.6–2.7).

The cross-paradigm boundary is diagnostic: covariance pooling is a band-power readout that discards the phase-locked evoked component on which ERP paradigms rely. A frozen-probe and scalp-pattern analysis confirms that the model reads paradigm-appropriate topographies exactly where that representation carries the signal. What generalizes is therefore the *signal family* rather than the named paradigm. Together these results establish geometry-conditioned hyperalignment as a practical route to neural decoders that are generalizable across datasets and across paradigms within that signal family.

## 2 Results

The results are organised around a single argument. We first show that the geometry-conditioned frontend places every montage (seen or unseen) into one shared anchor space (Sections 2.1–2.2). We then establish that this shared space is what makes decoding transfer across datasets (Sections 2.3–2.4). Next, we show that pairing the learned frontend with the Riemannian SPD backbone is what amplifies that transfer (Section 2.5). Finally, and most notably, the same geometry transfers across *paradigms* (Sections 2.6–2.7). The final subsection characterises robustness to the frontend’s design choices (Section 2.8).

### 2.1 The hTSMNet concept: from source–leadfield–sensor physics to a shared anchor space

#### The physical world

The activity we want to decode arises from a small set of latent neural sources (Figure 1, bottom plane). Volume conduction maps these sources to the scalp through a subject-specific *leadfield matrix* (the forward model *L_s_*), set by head geometry, tissue conductivity, and electrode placement. Denoting the *P* latent sources as **u**, a recording’s sensor signal is

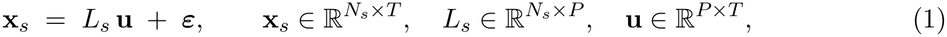

and is then sampled by that subject’s particular electrode montage (top plane). Two recordings of the same neural process therefore differ for two distinct reasons: their montages differ (channel count, layout, reference), and their leadfield matrices *L_s_* differ even for identical sources **u**. Therefore, cross-dataset transfer is not merely a distribution-shift problem: the recordings do not even share an observation space. This is why a shared encoder that is fed with raw channels cannot fix it: it must discard non-shared electrodes or learn a topology-agnostic map, and in neither case does it account for the underlying source-to-sensor geometry. (We give the forward model and all formal definitions in Methods, Section 4.1.)

#### The anchor space

hTSMNet inserts a single *shared* coordinate system between the (unknown) sources and the (montage-specific) sensors. We fix a number of *D anchor* positions on a canonical scalp—one template common to all subjects and datasets (Figure 1, middle plane)— and define the *anchor space* as the *D*-dimensional space of signals carried by those anchors. For each recording, a learned linear *mapping A_s_ ∈* ℝ*^N_s_×D^* reads its native sensors onto the anchors,

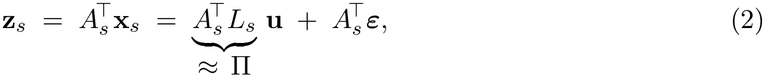

so that whatever the input montage, the downstream model always receives a fixed-*D* anchor signal **z***_s_*that lives in the same space for every recording (Figure 1). The net source-to-anchor map is the bracketed composite 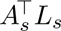, which the alignment must make consistent across recordings. Here Π is a *conceptual* target rather than a measured quantity, since *L_s_* is unknown and we never observe the composite directly (Section 3.2 returns to the evidence that training does drive it toward one shared operator).

#### What *A_s_* is—and what it is not

The alignment *A_s_* is a *learned, geometry-conditioned* sensor-to-anchor map—not a leadfield inverse and not a source-reconstruction filter (the object driven toward consistency is the composite 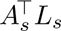, never the unknown leadfield matrix *L_s_* itself). Crucially, *A_s_* = *A*(*C_s_*) is produced from *geometry alone*, by one shared function of the electrode configuration *C_s_* with *no subject-specific parameters*. So Equation (2) is the zero-shot forward path: an unseen subject, dataset, or montage contributes only its coordinates, from which *A_s_* and the anchor signal **z***_s_* follow with no calibration—the basis of every zero-shot result below. We defer to the Discussion (Section 3.2) why reading the anchors as a shared coordinate frame, rather than recovered cortex, is the right view.

The aligned anchor signal is then forwarded to the TSMNet backbone [13], which pools its spatial covariance and classifies it on the symmetric positive-definite (SPD) manifold (Section 4.1). We chose a Riemannian SPD backbone, rather than a generic convolutional or transformer network, for reasons of physiological fit, sensitivity to the frontend, and interpretability that we return to in the Discussion (Section 3.4).

### 2.2 Constructing the alignment matrix: geometry plus a learned correction

Section 2.1 treated the alignment *A_s_*as a single map from sensors to anchors. We now decompose that map and show it is both learnable and geometrically consistent across subjects, the two properties the rest of the paper relies on. Intuitively, the frontend does two things: it first *interpolates* each recording’s sensors onto the anchors by physical proximity, a fixed geometric prior, and then *corrects* that interpolation with a task-selected learned adjustment. Concretely, the alignment combines (i) a radial-basis-function (RBF) interpolation kernel, which weights each sensor’s contribution to an anchor by their distance with a learned per-anchor width, and (ii) an additive correction produced by a single shared, position-conditioned network. Their sum is retracted onto a near-orthonormal projector so that the anchor axes stay non-redundant. Both parts are shared across all subjects and trained end-to-end with the classification loss(the full construction, including kernel, correction network, projection, and training is given in Section 4.1).

**Figure 1:**
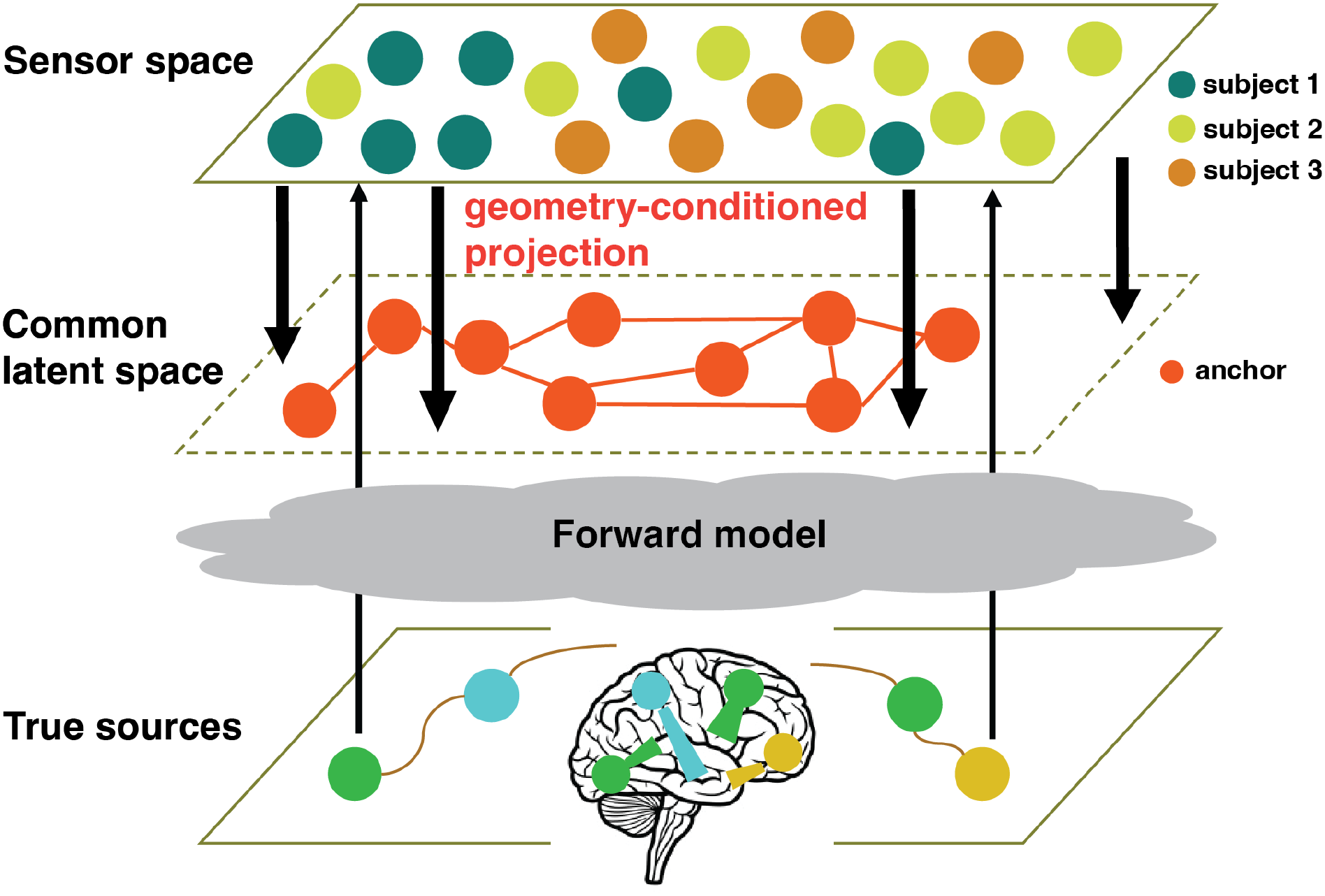
The hTSMNet concept: two maps, kept distinct. Latent neural sources (bottom plane, on the cortex) reach the scalp through the true, subject-specific *leadfield matrix*—the physical *forward model* (grey cloud)—and are sampled by each subject’s heterogeneous *sensor* montage (top plane; one colour per subject), so two recordings of the same neural process need not share an observation space. hTSMNet inserts a single shared *anchor space* of *D* fixed anchors (middle plane, orange; labelled “common latent space” in the schematic) and learns a separate, *geometry-conditioned alignment A_s_*(thick arrows) that maps each montage’s sensors onto those shared anchors. These anchors are reference scalp locations, not recovered sources. The forward model (sources *→* sensors) is the true physics and is unknown to us. The geometry-conditioned alignment (sensors *→* anchors) is the map hTSMNet controls—a learned surrogate for a shared source geometry, not an estimate of the physical inverse 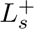 (Eqs. 1–2). The resulting anchorspace signal 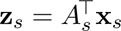 is then forwarded to the TSMNet backbone (Section 4.1).

Figure 2A–C visualises the two components and their composition for one source montage (Lee2019-MI, 56 channels). Because the alignment carries no per-subject parameters, this map is shared by every subject recorded on that montage, so the fold holds only *two* distinct source alignments, one per source dataset. The RBF base (A) has the expected sparse, local structure— each anchor draws from a small neighbourhood of sensors—while the learned correction (B) adds a broader, sign-varying pattern. The resulting alignment *A_s_* (C) departs substantially from the geometry-only RBF projector: the relative change—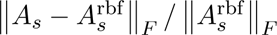, the Frobenius norm of the learned-minus-RBF difference normalised by the RBF norm—is 0.75 for this montage (projected-space cosine 0.74, *i.e.* the cosine between the vectorised *A_s_* and 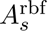) and 0.66 for the fold’s other source montage (PhysionetMotorImagery, 64 channels). Unlike a fixed interpolation rule, the learned alignment is free to re-weight which sensors feed each anchor. We show below that this freedom is what the downstream covariance representation needs.

**Figure 2:**
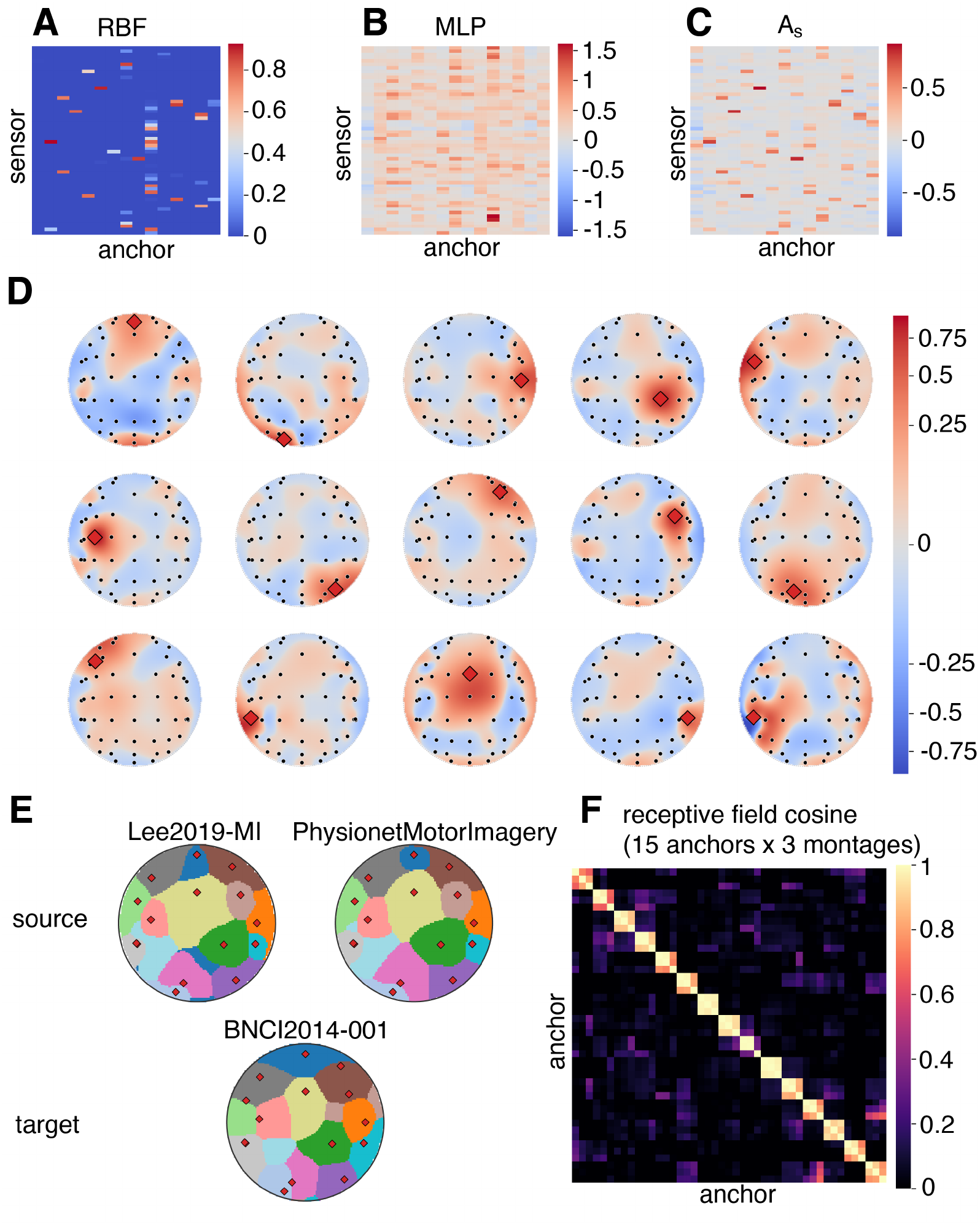
The geometry-conditioned alignment. *A_s_***. (A–C)** The two components and their composition for one source montage (Lee2019-MI; 56 channels, *D* = 15; sensor rows *×* anchor columns): the RBF base **(A)**, the learned correction **(B)**, and the retracted alignment *A_s_* **(C)**. The learned alignment departs from the geometry-only RBF projector by 0.75 in relative Frobenius norm for this montage (projected cosine 0.74) and 0.66 for the fold’s other source montage (PhysionetMotorImagery, 64 channels). **(D)** Receptive field of each of the *D* = 15 anchors for the same montage—the alignment column *A_s_*[:*, j*] interpolated on the scalp. The red diamond marks anchor *j*. Fields are centred on their anchor (locality preserved) yet form anisotropic spatial filters that recruit distant sensors. **(E)** Anchor parcellation each montage induces—each scalp location coloured by its dominant anchor, red diamonds mark anchor positions—for the two source montages and the zero-shot BNCI2014-001 target. The partitions coincide (source–zero-shot-target field cosine 0.80; 65% of target scalp pixels share the source partition), so the geometry-conditioned frontend places an unseen montage into the same anchor space. **(F)** Cosine matrix of all 45 anchor receptive fields (*D* = 15 anchors *×* 3 montages) on a common scalp grid. Bright 3 *×* 3 diagonal blocks (same anchor across montages, mean 0.85) and dark off-blocks (different anchor, 0.01) show that each anchor has a single, montage-independent, mutually-orthogonal receptive field. Because *A_s_* has no subject-specific parameters, these three montages are the only distinct alignments in the fold.

The learned alignment nonetheless remains geometrically interpretable. Figure 2D plots, for the same montage, the *receptive field* of each of the *D* = 15 anchors—the column *A_s_*[:*, j*] rendered on the scalp, with the anchor position marked by a red diamond. Every receptive field is spatially coherent and centred on or near its anchor, so locality is preserved. Yet most are not simple isotropic bumps: they extend lobes toward distant sensors and develop the positive/negative structure of a spatial filter, a behaviour a fixed RBF kernel cannot express. The frontend has thus learned, from the task alone, to combine local interpolation with longer-range spatial contrasts.

Finally, Figures 2E–F ask whether these receptive fields define a shared coordinate system or merely a per-montage artefact. Adding the zero-shot target to the two source montages above, the fold has just *three* distinct alignments in total: the two source datasets (Lee2019-MI, 56 channels; PhysionetMotorImagery, 64) and the held-out BNCI2014-001 (22 channels), the last aligned zero-shot from geometry alone. Figure 2E renders the anchor parcellation each montage induces, with each scalp location coloured by its dominant anchor. The partitions coincide across the two source montages and the zero-shot target (source–source receptive-field cosine 0.95, source–zero-shot-target 0.80; 87% and 65% of scalp pixels share the source partition), so the geometry-conditioned frontend carves an unseen montage into the same anchor territories the model was trained on. Figure 2F makes this quantitative: the cosine matrix of all 45 receptive fields (*D* = 15 anchors *×* 3 montages), interpolated on a common 24*×*24 scalp grid, has 3*×*3 diagonal blocks (one anchor across the three montages) averaging 0.85 against 0.01 for differentanchor entries. Each anchor thus maps to a single, montage-independent receptive field that is near-orthogonal to the others. Within a montage the *D* columns are orthonormal by construction (max *|*off-diagonal*|* of *A^⊤^A_s_ ≤* 0.003 for the dense source montages; the sparse 22-channel target reaches 0.32, where thin sensor coverage leaves the polar retraction only approximately orthonormal). The shared anchor geometry is what carries across recordings.

### 2.3 Cross-dataset transfer: the LODO few-shot benchmark

We have shown what the frontend builds—a shared anchor space that even unseen montages enter from geometry alone. We now evaluate that space, beginning with the question that motivated the architecture: does a decoder trained in this shared space transfer to an entirely held-out dataset?

#### Protocol

We adopt a leave-one-dataset-out (LODO) protocol: each method is trained on the union of the source datasets and evaluated on a held-out target dataset that was never seen during training. All methods decode the binary left-hand vs. right-hand (LH/RH) subtask, with *k ∈ {*0, 5, 10, 20, 50*}* labelled support (calibration) trials per class used to adapt only the final-layer classifier (Section 4.3). The *k* = 0 setting is strict zero-shot. The three folds hold out BNCI2014-001 (22 channels, 9 subjects), Lee2019-MI (56 channels, 54 subjects), and PhysionetMotorImagery (64 channels, 109 subjects). Pooled over folds the corpus spans 172 subjects. Unless stated otherwise, every accuracy in this paper is *balanced accuracy* and chance is 50%. Every entry is the mean of the per-target-subject mean accuracy across five independent random seeds (controlling network initialisation and data ordering). For its headline configuration, hTSMNet uses a *data-agnostic* anchor template (*D* = 15 farthest-point samples of a canonical `standard_1020` layout using canonical standard 10–20/10–05 electrode positions looked up from channel labels via MNE, then reduced to the top-down *x*–*y* projection), so the main result requires no paradigm-specific tuning. To compare fairly across heterogeneous montages, the classical and specialist comparators—which cannot ingest a variable montage—are first mapped onto the same *D* = 15 canonical anchors by a *fixed, unlearned* RBF interpolation (denoted RBF(*D*=15) in Table 3): a single global kernel width and no polar retraction—that is, the raw RBF *base* of our frontend with *neither* its learned per-anchor widths *nor* its polar step, and hence not identical to hTSMNet’s learned frontend or even its geometry-only ablation (Section 4.3). The foundation models instead keep their own pretraining montages. Handing every fixed-input comparator this shared coordinate system is intentionally generous—it is what lets them decode across heterogeneous montages at all—so the remaining comparison isolates what is gained by *learning* the alignment and coupling it to the SPD backbone (full training details are in Methods, Section 4.1).

#### Zero-shot is competitive—the advantage emerges with calibration

Figure 3A–C shows balanced accuracy versus *k* for hTSMNet and the eight comparators. At strict zeroshot (*k* = 0), hTSMNet attains 68.8 *±* 3.7%, 64.5 *±* 1.7%, and 68.5 *±* 1.8% on BNCI2014-001, Lee2019-MI, and PhysionetMotorImagery (three-fold mean 67.3 *±* 1.7%). This is marginally ahead of the best single comparator (Deformer, 66.4%) but without a clear per-dataset edge: hTSMNet leads EEGNet on BNCI2014-001 by only 0.6 pp, sits within seed noise of Deformer on Lee2019-MI, and trails the channel-matched foundation model NeuroGPT on PhysionetMotorImagery (68.5 vs. 71.4%). The advantage becomes clear as soon as a handful of labelled trials are available. At *k* = 20, hTSMNet reaches 80.7 *±* 0.6%, 70.1 *±* 0.6%, and 76.3 *±* 0.9% (three-fold mean 75.7 *±* 0.2%), ahead of Deformer—the strongest comparator at every fold (77.7, 65.9, 73.5%)—by 3.0, 4.1, and 2.8 pp, respectively (+3.3 pp on the three-fold mean). The lead persists at *k* = 50 (83.2 and 72.9% on the two folds where it is defined—PhysionetMotorImagery has no *k* = 50 budget—+3.2 and +4.6 pp over Deformer). The gap over the best comparator stays in a +2.5 to +3.9 pp band at every calibrated budget *k ≥* 5 (three-fold mean for *k ≤* 20, two-fold at *k* = 50), so it is not an artefact of any single shot budget.

**Figure 3:**
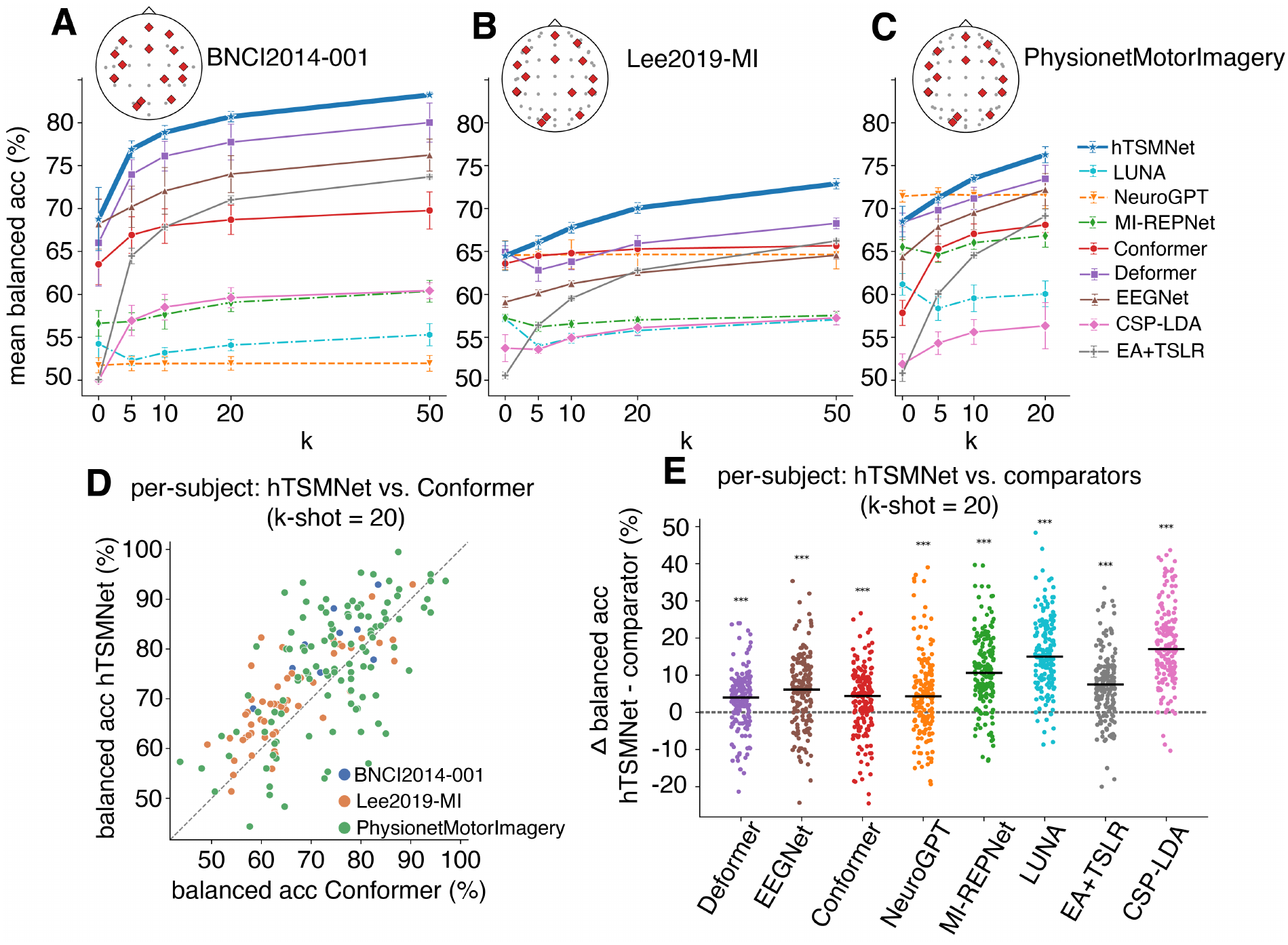
Cross-dataset LODO few-shot benchmark (LH vs. RH). (A–C) Mean balanced accuracy vs. *k*-shot per class for hTSMNet and eight comparators on the three held-out datasets. Topoplot insets show each target’s native montage with the 15 canonical anchors (red diamonds). Markers are the mean of the per-target-subject mean accuracy over five seeds. Error bars are the seed standard deviation. hTSMNet (blue) is the highest method for every *k ≥* 5 on all three datasets. At strict zero-shot (*k* = 0) it lies within the leading group (see text). **(D)** Per-subject paired scatter at *k* = 20 against the closest per-subject competitor (Conformer). Each point is one subject (coloured by dataset), and points above the diagonal are hTSMNet wins. **(E)** Per-subject advantage Δ = hTSMNet *−* comparator (pp) at *k* = 20 for each comparator. Black bars are medians, annotations give the win rate and Wilcoxon significance (Holm-corrected). In all eight comparisons hTSMNet is significantly more accurate at *p <* 10*^−^*^3^.

#### Baseline models

We consider three families of comparators: classical Riemannian pipelines, specialist deep networks, and EEG foundation models. *Classical Riemannian pipelines* (CSP– LDA, common spatial patterns with shrinkage LDA; EA–TSLR, Euclidean alignment with tangent-space logistic regression) sit near chance at *k* = 0 (50.6–51.9% three-fold mean): with no cross-dataset pretrained model they only begin to decode once support trials arrive (EA– TSLR reaches 71.0% on BNCI2014-001 at *k* = 20, still 9.7 pp below hTSMNet). *Specialist deep networks* (EEGNet, Conformer, Deformer) benefit from source pretraining and adapt steadily. Deformer is the strongest comparator overall, but never overtakes hTSMNet on any dataset once a few trials are available. *Pretrained foundation models* are mixed: NeuroGPT, constrained to its 22-channel template, is in fact the single best zero-shot method on PhysionetMotorImagery (71.4%), but its fixed template mismatches the other montages and it barely improves with *k*. The variable-montage model LUNA, evaluated on native montages, is the weakest pretrained network, reaching only 54–61% at *k* = 0. Only hTSMNet combines explicit geometric alignment with SPD covariance processing, and only hTSMNet leads across all three montages once calibrated.

#### The advantage holds subject by subject

Group means can hide heterogeneity, so we paired hTSMNet against each comparator *on the same subject* at the *k* = 20 operating point and tested the per-subject difference (Figure 3D–E; 171 of the 172 subjects, pooled across folds—the one excluded PhysionetMotorImagery subject has no *k*=20 score (Section 4.2)). hTSMNet wins on the majority of subjects against every comparator, and a one-sided Wilcoxon signed-rank test (Holm-corrected across the eight comparators) is significant at *p <* 10*^−^*^3^ in all eight cases. The smallest margins are against the strongest specialist deep networks—Conformer (70% of subjects, median Δ = +4.4 pp, *p* = 4 *×* 10*^−^*^7^; Figure 3D) and Deformer (72%, +4.0 pp, *p* = 2 *×* 10*^−^*^8^)—followed by NeuroGPT (68%, +4.3 pp) and EEGNet (73%, +6.2 pp). Against EA–TSLR hTSMNet wins 81% of subjects (+7.5 pp) and against MIRepNet 86% (+10.6 pp); against LUNA and CSP–LDA it wins 96% (+15 to +17 pp). The few-shot cross-dataset advantage is therefore robust at the level of individual subjects, not just dataset averages.

### 2.4 Why hTSMNet transfers across datasets: representational geometry

What property of the learned representation produces this cross-dataset advantage? A natural hypothesis is that hTSMNet simply compensates for the distribution shift between datasets, minimising their divergence in feature space. We find the mechanism is more specific, and is a property of the class direction rather than of dataset overlap. hTSMNet makes the left-/righthand discriminative direction point the same way in every dataset, so that a single linear decision rule learned on the source datasets is already oriented correctly on a held-out one. We quantify this on the LODO folds with the *class-direction consistency* CC: the mean pairwise cosine between the per-dataset LH/RH axes, computed on the penultimate (LogEig tangent-space) features and averaged over the three folds (Section 4.4).

Figure 4A,B makes the effect concrete on the BNCI2014-001 fold by plotting every trial against the frozen *k* = 0 class axis (*x*) and the dataset-centroid axis (*y*). Under no learning (raw EEG projected through geometry only, then covariance pooling and LogEig; panel A), the heldout dataset forms a compact cloud displaced far along the dataset axis from the source data, and its LH/RH marginal (bottom) is unseparated: the class axes of the three datasets barely agree (CC = 0.20). After hTSMNet alignment (panel B), the held-out dataset is drawn back into the source distribution and its LH/RH marginal separates along the shared class axis (CC = 0.69). Therefore, the class structure, not the dataset identity, now organises the representation.

**Figure 4:**
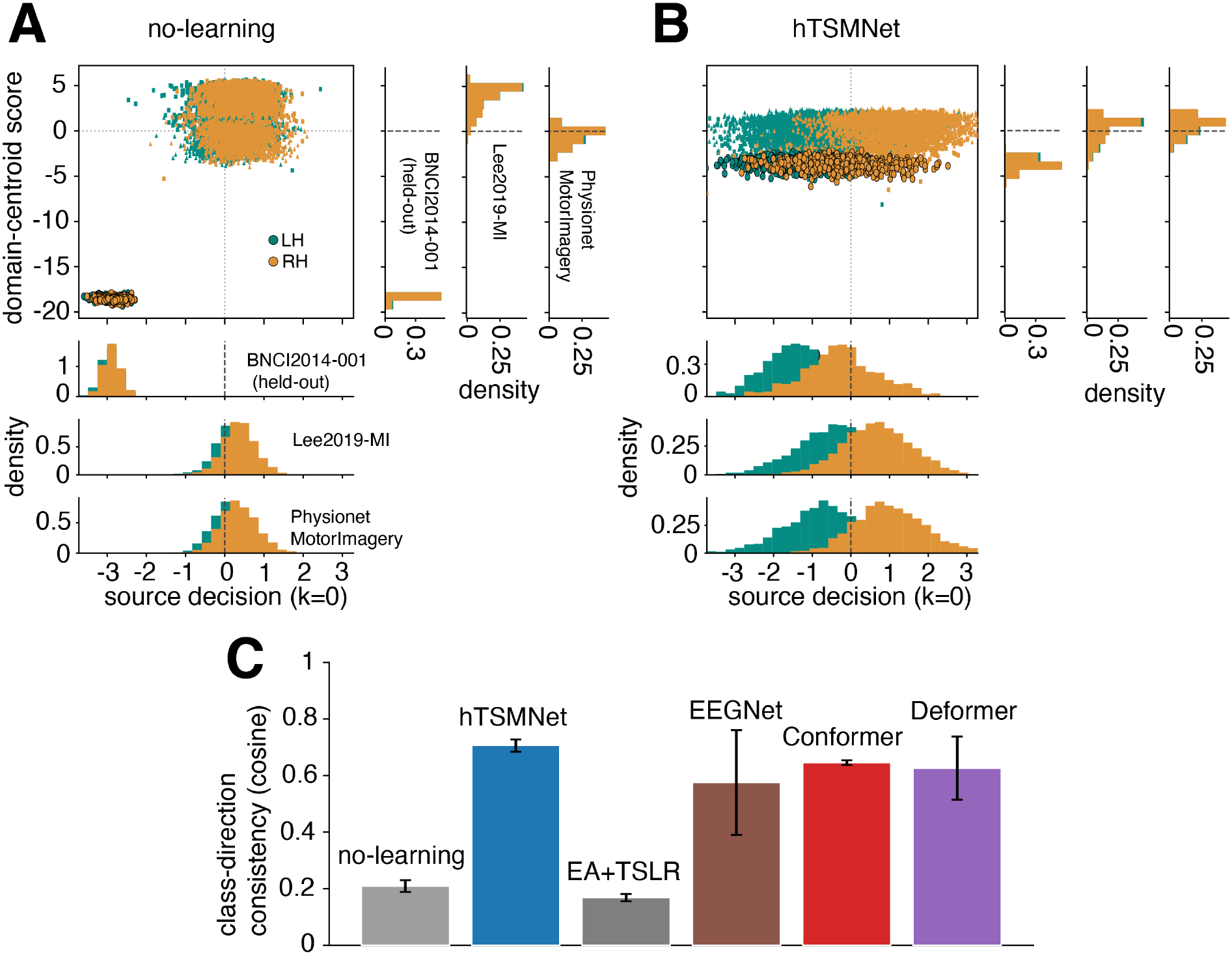
Cross-dataset representational geometry on the LODO folds. (A,B) Every trial on the BNCI2014-001 fold plotted against the frozen *k* = 0 class axis (*x*) and the domaincentroid axis (*y*). The held-out dataset is outlined. Marginals show each dataset separately: the bottom strips give its LH/RH split along the class axis (*x*), and the right strips its distribution along the domain-centroid axis (*y*). **(A)** No learning: the held-out dataset is displaced from the source datasets along the domain axis (right marginal), and its LH/RH split overlaps on the class axis (bottom marginal; class-direction consistency CC = 0.20). **(B)** hTSMNet: the held-out dataset is drawn into the source distribution along the domain axis (right marginal), and its LH/RH split separates along the shared class axis (bottom marginal; CC = 0.69). Colours: LH/RH trials are coloured teal and orange, respectively. **(C)** Cross-dataset classdirection consistency—the mean pairwise cosine of the per-dataset LH/RH axes (higher is better; mean*±*std over three folds). hTSMNet is highest at 0.71, versus 0.21 (no learning), 0.17 (EA– TSLR), and 0.58–0.64 for the trained deep comparators (EEGNet, Conformer, Deformer).

This pattern holds across folds (Figure 4C). hTSMNet attains the highest class-direction consistency of any method, CC = 0.71*±*0.02 (mean*±*std over the three folds)—about 3.4*×* the nolearning baseline (0.21*±*0.02) and well above the classical EA–TSLR pipeline (0.17*±*0.01), which does not align the class direction at all. The trained deep comparators do recover a partially shared class axis through end-to-end class supervision on the pooled source data (Conformer 0.64 *±* 0.01, Deformer 0.63 *±* 0.11, EEGNet 0.58 *±* 0.19), but none matches hTSMNet. This is what the structural anchor-space hypothesis predicts: every montage is mapped into one common anchor space rather than aligned dataset-by-dataset through class tuning. We note two caveats though. First, hTSMNet training also includes a mild *class-agnostic* covariancealignment regulariser (Section 4.1) that promotes cross-dataset distributional overlap but does not supervise the LH/RH direction itself. Second, the present analysis is correlational rather than a direct ablation of the anchor space: a direct test would randomise the sensor-to-anchor assignment and check that CC collapses, which we leave to a controlled follow-up. A class direction that is already shared across datasets is what makes hTSMNet’s few-shot calibration so efficient (Section 2.3): a classifier trained on the source datasets lands in nearly the right orientation on a new one, so only a handful of labelled trials are needed to close the gap.

### 2.5 The SPD backbone amplifies the learned frontend

The cross-dataset advantage raises a mechanistic question: does learning the alignment help on its own, or only when it is paired with the Riemannian SPD backbone? We answer it with a controlled transplant. The *same frontend architecture* is placed in front of three backbones: the deep SPD TSMNet, a shallow tangent-space SPD classifier, and a non-SPD convolutional network (EEGNet). Each frontend–backbone pair is trained *jointly end-to-end*, so a fresh frontend co-adapts to its own decoder (we do *not* freeze hTSMNet’s learned alignment and reuse its output as a fixed input). For each pair we measure the zero-shot gain from learning the alignment over the geometry-only RBF projector (Figure 5A). The gain scales with how geometric the decoder is: it is largest on the deep SPD backbone (+10.4 pp), smaller on the shallow SPD classifier (+4.6 pp), and smallest on the non-SPD convolutional backbone (+3.3 pp). The deep SPD backbone also reaches the highest absolute accuracy, and the benefit persists across *k* (Figure 5B). The learned frontend thus provides the largest benefit when coupled to a Riemannian decoder.

**Figure 5:**
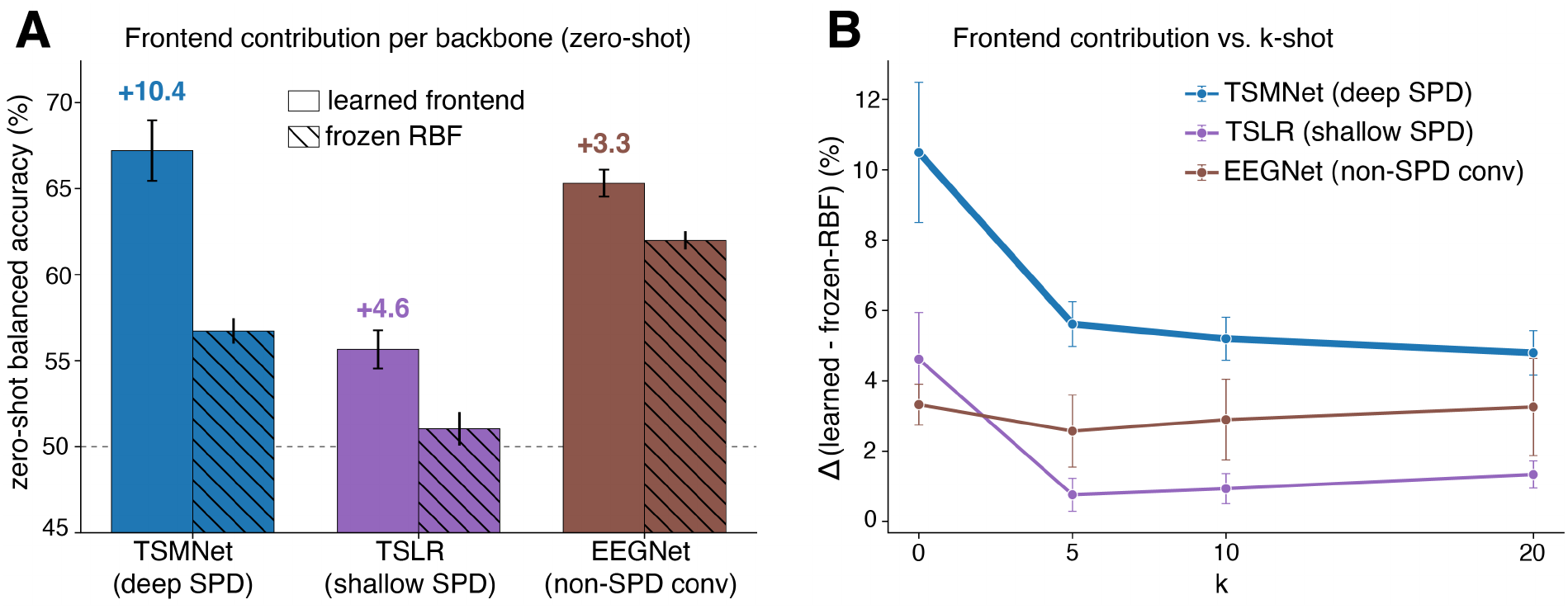
The learned frontend benefits a Riemannian backbone most. The same frontend architecture is paired with each of three backbones and trained jointly end-to-end with it (a fresh frontend per backbone, not hTSMNet’s learned alignment frozen and reused as input). **(A)** Zero-shot balanced accuracy with the learned frontend vs. the geometry-only RBF projector: the benefit of learning the alignment is largest on the deep SPD backbone (TSMNet, +10.4 pp), smaller on a shallow tangent-space classifier (+4.6 pp), and smallest on a non-SPD convolutional backbone (EEGNet, +3.3 pp). The deep SPD backbone also reaches the highest absolute accuracy. **(B)** The same benefit across *k*-shot.

### 2.6 Cross-paradigm transfer: a frozen MI model decodes other paradigms

The cross-dataset advantage rests on a stable shared class direction (Section 2.4) that the SPD backbone amplifies (Section 2.5). We therefore asked whether that same geometry transfers not just across datasets of the same paradigm, but across *paradigms*. We pretrained hTSMNet once on the pooled three-dataset MI corpus (LH/RH), *froze the entire network* and transferred it to five target datasets from three unseen paradigms by adapting only linear classifier on *k* labelled support trials per class. The MI-pretrained feature extractor is kept fixed throughout— the geometry frontend, temporal–spatial filters, covariance pooling, BiMap/ReEig layers, SPD batch-norm statistics, and LogEig tangent map—so this is a purely inductive test of the frozen representation. Only the tangent-space logistic final-layer classifier is re-fit, so successful transfer means that the MI-trained tangent representation already linearly encodes the new paradigm. The final-layer classifier is fit separately for each target subject: for each subject and support seed, we draw *k* support trials per class, fit a fresh classifier, and score that subject’s remaining trials. The frozen feature extractor is shared across subjects, no session split is used, and the five support draws define the reported seed average. The targets are mental-workload estimation (Hinss2021, 15 subjects, 61 channels), SSVEP decoding (Lee2019-SSVEP, 54 subjects; Kalunga2016, 12 subjects, 8 channels), and P300/ERP detection (BrainInvaders2013a, 24 subjects; BNCI2014008, 8 subjects). All identical comparators are transferred the same way (frozen body, classifieronly adaptation). Results are averaged over five adaptation seeds (Figure 6). Because each new paradigm needs its own output classifier, the *k*=0 point is at chance by construction. The meaningful question is the rate and extent of improvement under few-shot adaptation.

**Figure 6:**
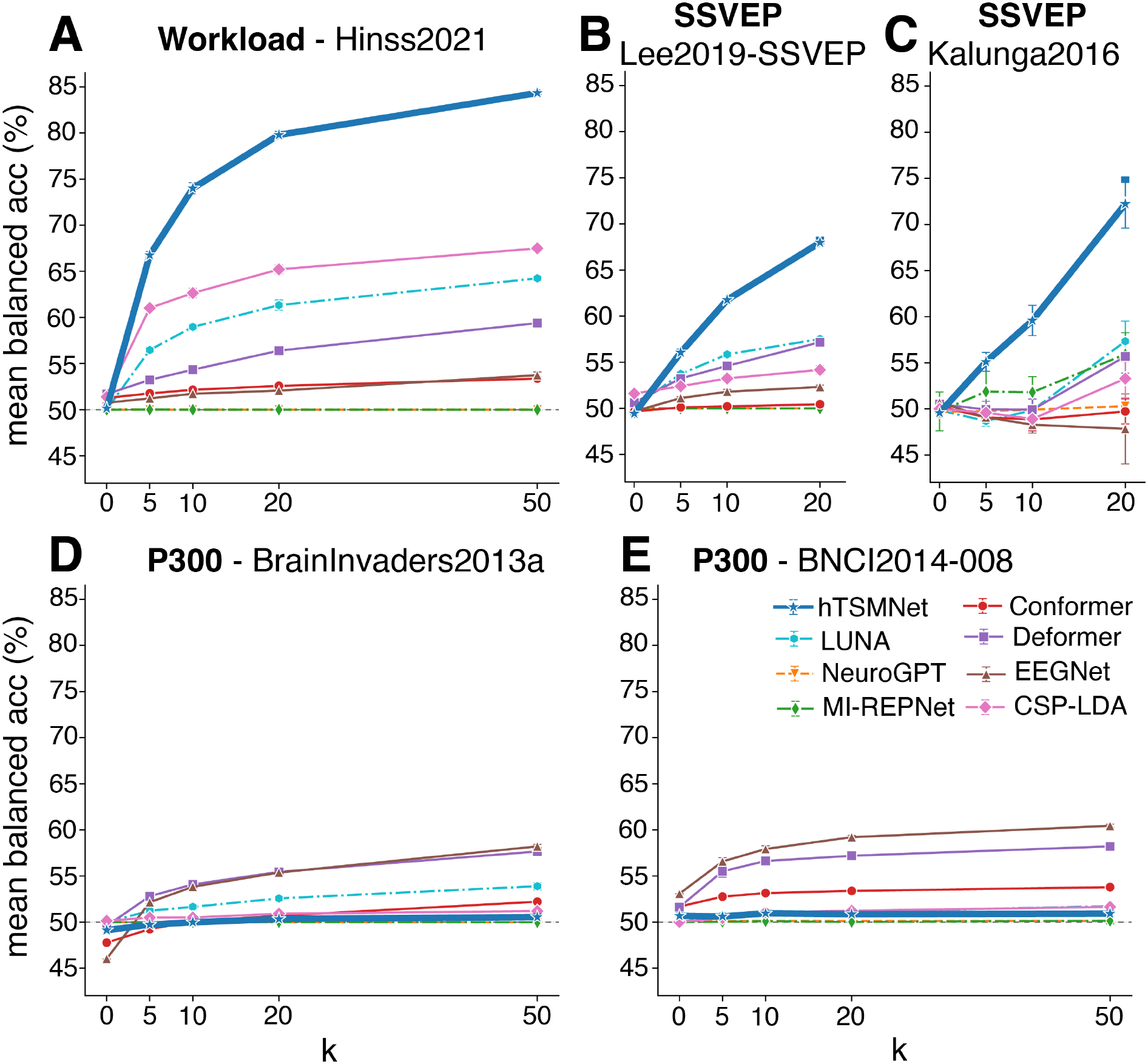
Cross-paradigm transfer from a frozen MI-pretrained model. Each method is frozen and transferred to an unseen paradigm by re-fitting only a linear classifier on *k* support trials per class (mean*±*std over five seeds; dashed line = chance). hTSMNet and the non-foundation comparators are MI-pretrained, while the foundation models use their official pretrained checkpoints. **(A)** Workload (Hinss2021): hTSMNet (blue) climbs to 84.3% at *k*=50, far above every other frozen-pretrained model. **(B–C)** SSVEP (Lee2019-SSVEP, Kalunga2016): hTSMNet reaches 68–72% at *k*=20. **(D–E)** P300 (BrainInvaders2013a, BNCI2014-008): hTSMNet (and the other covariance/band-power methods) remain at chance, while time-domain networks (EEGNet, Deformer) transfer. EA–TSLR is omitted as a within-target re-fit upper reference rather than a transfer method (see text).

#### Strong transfer to oscillatory paradigms

On mental workload, the frozen MI hTSMNet climbs from chance at *k* = 0 to 66.8% (*k*=5), 74.0% (*k*=10), 79.8% (*k*=20), and 84.3% (*k*=50)—a +34 pp gain over chance driven entirely by re-fitting a linear classifier on a frozen MI representation (Figure 6A). On SSVEP it reaches 68.0% (Lee2019-SSVEP, *k*=20) and 72.2% (Kalunga2016, *k*=20; Figure 6B–C). Among the frozen pretrained networks hTSMNet is the only one that transfers substantially: on workload at *k*=50 the next-best pretrained model is LUNA at 64.2%, followed by Deformer (59.4%), EEGNet (53.7%), and Conformer (53.4%), while NeuroGPT and MIRepNet stay at chance. The from-scratch CSP–LDA baseline, which re-fits entirely on the target, reaches 67.5%—*i.e.* with no feature-extractor updates (only a re-fit linear classifier), hTSMNet’s frozen MI features outperform a covariance pipeline trained from scratch on the workload data. For reference, a within-target *upper reference* (EA–TSLR, which re-estimates its full alignment and decoder on the target and is therefore not a transfer method; omitted from Figure 6) reaches 91.3% on workload and 73.3% on Lee2019-SSVEP at the same budgets—an approximate upper marker, not a strict bound—confirming that the oscillatory signal is clearly present and that hTSMNet recovers most of it from frozen MI features alone.

#### No transfer to evoked (ERP) paradigms

On both P300 datasets the frozen hTSMNet stays at chance regardless of *k* (50.7% *→* 50.9% on BNCI2014-008 and 49.1% *→* 50.6% on BrainInvaders2013a from *k*=0 to *k*=50; Figure 6D–E)—and so do the other covariance/bandpower methods (CSP–LDA, EA–TSLR). The *time-domain* networks behave oppositely: EEGNet and Deformer, whose convolutions retain the trial-aligned waveform, transfer to P300 (58–60% at *k*=20–50) far better than they do to the oscillatory paradigms. This double dissociation— covariance methods win on oscillatory paradigms and lose on ERP, time-domain methods do the reverse—is not an unexplained shortcoming but a direct prediction of what an MI covariance representation should and should not carry. We test that prediction directly in the next section.

### 2.7 The representational basis of cross-paradigm transfer

The cross-paradigm pattern in Figure 6 follows from a single property of the architecture: hTSMNet’s backbone pools *covariance*, which is a band-power (oscillatory) summary that discards phase-locked structure. We substantiate this with three frozen-representation diagnostics (Figure 7A–C) and a scalp-pattern traceback (Figure 7D–G), all computed on the frozen MI model. First, is the task signal present in the frozen features at all? We trained a linear probe on the frozen MI latent for each target (Figure 7A). The probe is well above chance for every oscillatory target—0.97 (Hinss2021 workload), 0.81 (Lee2019-SSVEP), 0.73 (Kalunga2016)—and for the MI references (0.76–0.88), but sits exactly at chance for both ERP targets (0.50, 0.50). The frozen MI representation contains an oscillatory readout of these tasks but not an ERP readout. The classifier-adaptation result of the previous section is the operational consequence.

**Figure 7:**
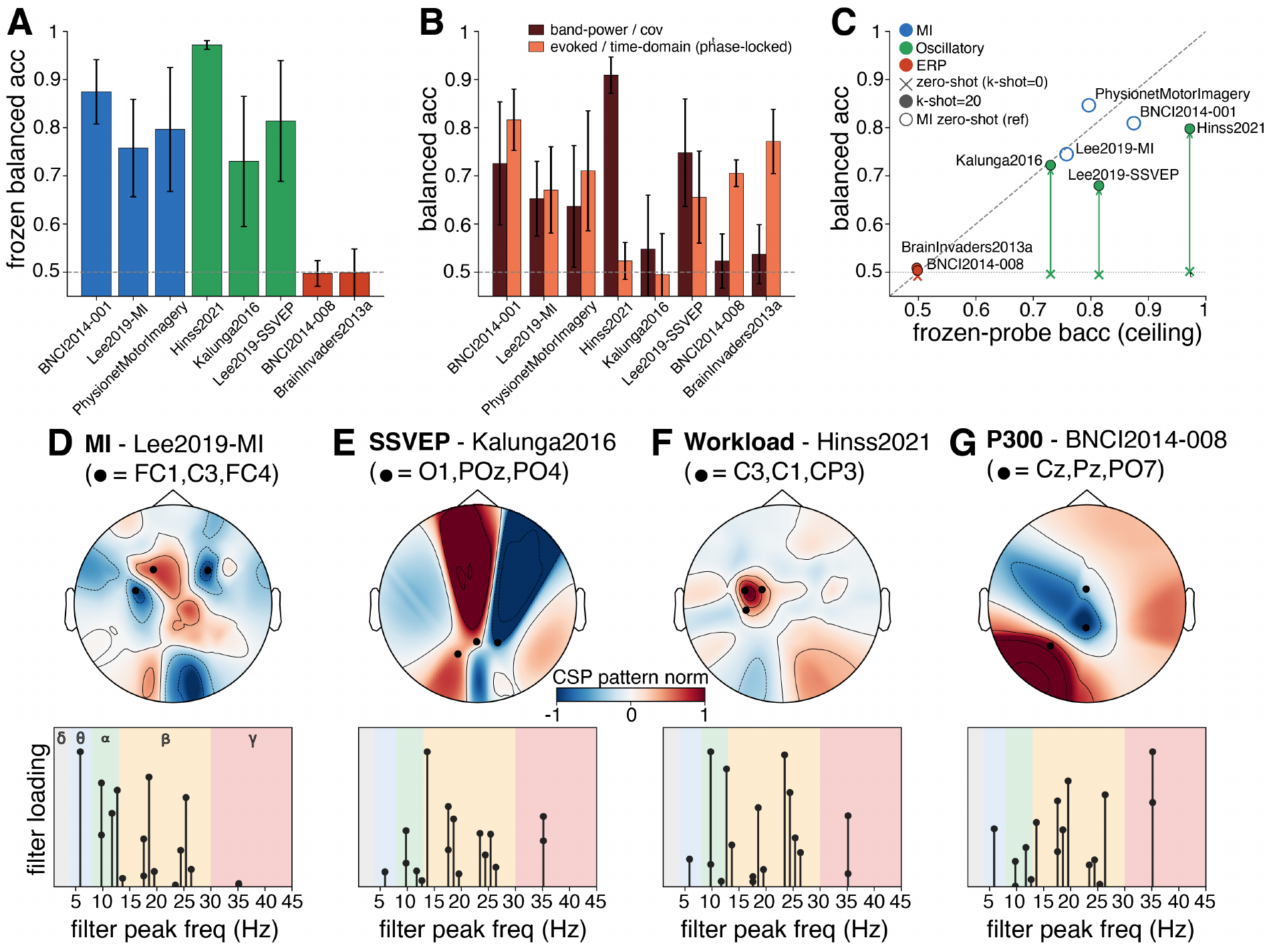
hTSMNet transfers along the band-power manifold. All panels use the frozen MI-pretrained model. **(A)** Frozen-probe balanced accuracy (“is the signal present?”) per target: high for MI (blue) and oscillatory (green) paradigms, at chance for ERP (red). **(B)** Class decoding from band-power/covariance features (what hTSMNet pools) vs. evoked/time-domain features (phase-locked): band-power dominates for oscillatory paradigms, evoked dominates for ERP—covariance pooling is largely insensitive to the phase-locked component. **(C)** The probe ceiling (*x*) tracks achievable transfer: *k*=0 (x) climbs to *k*=20 (filled) toward the ceiling for oscillatory targets, while ERP targets stay pinned at chance (MI zero-shot references shown as open circles). **(D–G)** Dominant decoding direction traced through the frozen frontend onto the scalp (sensor-space topographies, not cortical source estimates; top-3 sensors annotated). The per-filter spectral stem below each topography places every learned temporal filter at its peak frequency (horizontal axis), with stem height *|α_ft_|* giving that filter’s loading on the dominant CSP source (unnormalised relative scale): sensorimotor for MI (D), occipital for SSVEP (E), central for workload (F), and centro-parietal for ERP (G; correct P300 topography but no decodable band-power signal).

Second, why is the ERP signal absent? For each target we decoded the class label from two feature views of the same trials: band-power/covariance features (what hTSMNet’s pooling sees) versus evoked/time-domain features (the trial-averaged, phase-locked waveform). The two views dissociate exactly along paradigm lines (Figure 7B). For oscillatory targets the band-power view dominates (workload 0.91 vs. 0.52 evoked; SSVEP 0.75 vs. 0.66). For ERP targets the ordering reverses sharply (BNCI2014-008 0.52 band-power vs. 0.71 evoked; BrainInvaders2013a 0.54 vs. 0.77). The P300 signal is concentrated in the phase-locked component that covariance pooling integrates away, so classifier adaptation alone cannot recover it.

Third, these two diagnostics combine into a quantitative relationship: the frozen-probe accuracy acts as an empirical *ceiling* that tracks how far few-shot adaptation can climb (Figure 7C). Plotting each target’s *k*=20 transfer accuracy against its probe ceiling, the oscillatory targets rise from chance at *k*=0 toward their (high) ceilings, the MI references sit near the diagonal, and the ERP targets remain near chance on both axes, where the ceiling itself is at chance. Across these five targets transferability is thus not idiosyncratic: it is ordered by whether the paradigm’s discriminative signal lives on the oscillatory band-power manifold the frozen model represents.

Finally, when the signal is present, does the frozen model read it from the expected scalp regions? To recover the spatial pattern we fit common spatial patterns (CSP) on the frozen MI latent of each paradigm, take its more spatially focal dominant component, and back-project that anchor-space direction through the frozen frontend onto the scalp (Figure 7D–G; these are sensor-space topographies, not cortical source estimates; full procedure in Section 4.5). The recovered scalp pattern matches the known neurophysiology of each paradigm. The MI pattern is sensorimotor and lateralised (top sensors FC4, C3, FC1; Figure 7D), consistent with mu/beta event-related desynchronisation over motor cortex [16]. The SSVEP pattern is occipito-parietal (PO4, O1, POz; Figure 7E), consistent with visual-cortex steady-state generators [17]. The workload pattern is central/centro-parietal (C3, C1, CP3; Figure 7F), consistent with sensorimotor-/parietal-distributed workload modulation [18]. The ERP pattern falls on centro-parietal sites (Pz, PO7, Cz; Figure 7G), the canonical P300 topography [19]—but here the spatial pattern is correct while the decodable signal is absent, precisely because the diagnostic value of P300 is carried by phase-locked timing rather than by band power. For the spectral view we place each learned temporal filter at its peak frequency, weighted by its loading on that same decoding direction. These loadings cluster in the 6–35 Hz range spanning *θ*/*α*/*β*, the bands that carry oscillatory task information. Taken together, Figure 7 shows that hTSMNet transfers exactly where its band-power representation carries the signal, recovers the correct scalp topography where it does, and fails where the signal is phase-locked.

### 2.8 Ablations and robustness

Three further analyses characterise how robust the cross-dataset results are to the frontend’s two design choices (the anchor count and the anchor template) and to one optional element of the evaluation protocol, test-time adaptation. Each is detailed in a supplementary figure.

#### Sensitivity to the anchor count

1. *D*. The anchor dimension *D* is the frontend’s main hyperparameter. Sweeping *D ∈ {*5*, …,* 60*}* at zero-shot (Figure S1) shows that hTSMNet’s three-foldmean accuracy rises steeply from *D*=5 (59.9%) to *D*=10–15 (66.9–67.3%) and then *saturates*, holding a *∼* 64–68% plateau out to *D*=60. The same saturation is visible for the specialist deep networks—a denser RBF-projected input gives their convolutions finer spatial resolution, with diminishing returns past *D≈*15—and at zero-shot hTSMNet is competitive with the strongest of them across the sweep rather than uniformly ahead (consistent with Section 2.3). Two practical points follow. First, *D*=15 sits at the knee of the curve and is our default. Second, we keep *D ≤* min*_s_ N_s_* across the corpus as a conservative guideline: once *D* exceeds a dataset’s channel count the aligned covariance is rank-deficient (and the subsequent BiMap layer lowers its effective rank further), so its LogEig tangent representation departs from the full-rank case—the regime the 8-channel cross-paradigm targets fall into (Section 4.2). On the 22-channel BNCI2014-001 this guideline is precautionary rather than forced by the data: the zero-shot curve is noisy once it saturates, its one salient dip sits at *D*=20 (still full-rank, *D < N_s_*) and is driven by a subset of seeds collapsing toward chance rather than by rank, and we see no systematic degradation deeper in the rank-deficient regime. We therefore treat *D ≤* min*_s_ N_s_* as a deployment guideline, not an architectural limit.

#### Robustness to the anchor template

Because the anchors define the shared coordinate system, one might worry that results hinge on placing them well. They do not (Figure S2). At *D*=15, three template constructions—the data-agnostic canonical (farthest-point) default, *k*-means of the pooled sensor cloud, and a random draw—land within *∼* 1.5 pp of one another at every *k*, confirming that hTSMNet is insensitive to the exact anchor placement. A paradigminformed template that pins anchors over the sensorimotor strip does help, but modestly: it adds *∼* 1.9 pp at zero-shot on the three-fold mean (up to +4.3 pp on BNCI2014-001, 73.1 vs. 68.8%) and +1.2 pp at *k*=20. We deliberately report the *data-agnostic* template as the headline configuration: it requires no domain knowledge of the paradigm, matches the template shared with the comparators and the *D*-sweep, and shows that hTSMNet’s advantage does not depend on hand-tuned anchors. In practice, choose anchors by farthest-point sampling on the canonical `standard_1020` scalp, set *D ≈* 15 at the accuracy knee, and keep *D ≤* min*_s_ N_s_*whenever possible to avoid low-rank covariances. A paradigm-specific template is then an optional, small improvement when prior knowledge of the neural substrate is available. In this sense a paradigminformed template acts as a *brain-state-aware* prior—it spends anchor capacity on the cortex a paradigm is known to engage—and the modest gain (*∼* 1.9 pp zero-shot) quantifies how much such prior knowledge is worth on top of the data-agnostic default.

#### Test-time adaptation and the zero-shot ceiling

Throughout the main comparison every method is evaluated *without* test-time adaptation, so that hTSMNet is compared under the same evaluation settings as the non-BatchNorm baselines (CSP–LDA, EA–TSLR, LUNA), and *k*=0 is strict zero-shot (no target labels, no target statistics). But *test-time AdaBN* —recomputing a network’s BatchNorm running statistics from the held-out subject’s own *unlabelled* trials— is available to any BatchNorm architecture, costs no labels and no weight updates, and is computed once per subject at inference. Because it uses the target’s unlabelled trials, it is best understood as a *transductive* (label-free) zero-shot step rather than as strict zero-shot. Crossdataset transfer is exactly the setting where it should matter, because the held-out montage and subject distribution differ from the training pool. Yet LODO comparisons in the literature rarely grant it to the comparators. We therefore re-evaluated the four BatchNorm decoders (hTSMNet, Deformer, EEGNet, Conformer) with AdaBN on, at zero-shot (Figure S3).

Two findings follow. First, AdaBN is the single largest lever on zero-shot LODO: recomputing BatchNorm on the target lifts every decoder—hTSMNet +4.7, Conformer +4.8, Deformer +3.8, EEGNet +2.2 pp (three-fold mean)—so a comparison that withholds it from the baselines understates them substantially (Deformer, for instance, rises from 66.4 to 70.2%). Second, the ranking is unchanged: even on this fairest BatchNorm-vs-BatchNorm footing hTSMNet remains the best zero-shot method (71.9 *±* 0.5%), ahead of Deformer (70.2 *±* 0.7), Conformer (66.5 *±* 0.3) and EEGNet (66.1 *±* 1.2), and it leads on all three held-out datasets (BNCI2014-001 76.1 vs. 74.7; Lee2019-MI 67.3 vs. 65.5; PhysionetMotorImagery 72.5 vs. 70.5). hTSMNet’s zero-shot edge—+1.7 pp over the strongest comparator—is thus present but modest, not an artefact of who was allowed test-time adaptation.

For hTSMNet the AdaBN gain is confined to zero-shot: it collapses to +0.6 pp by *k*=5 and +0.2 pp by *k*=20, because a handful of labelled calibration trials already supply the same subject-specific recentring. Test-time AdaBN is therefore an optional, unsupervised boost for the no-calibration regime. It does not alter the paper’s central claims, which rest on the AdaBNindependent few-shot lead (Section 2.3) and on cross-paradigm transfer (Sections 2.6–2.7).

#### Summary of contributing components

Table 1 collects the ablations above into a single progression, comparing hTSMNet with the strongest comparator (Deformer) on the three-foldmean LODO MI benchmark as geometric and adaptation components are added in turn. Three points stand out. First, the comparator is *already* granted the hTSMNet geometric coordinate system: with no frontend learning at all, a fixed RBF projection onto the canonical anchors lets Deformer decode across the heterogeneous montages (66.4%), and at this geometry-only stage it is in fact ahead of the geometry-only input fed to hTSMNet’s SPD backbone (56.9%). Second, hTSMNet’s gains come from *learning* the alignment and from the SPD backbone: adding the learned per-anchor RBF width (64.3%, +7.4 pp) and then the learned MLP correction (67.3%, a further +3.0 pp) lifts the geometry-only input by +10.4 pp at zero-shot—most of it from learning the per-anchor scale—and these learned components have no analogue for a fixed-input network (“n/a”). Third, both adaptation levers preserve hTSMNet’s lead: few-shot classifieronly calibration (*k*=20) reaches 75.7% versus Deformer’s 72.4% (+3.3 pp), and label-free testtime AdaBN reaches 71.9% versus 70.2% (+1.7 pp). The learned frontend and the SPD backbone, not the shared geometric input, are what separate hTSMNet from the strongest comparator.

**Table 1:** Summary ablation: contribution of each component. Three-fold-mean balanced accuracy (%) on the cross-dataset LODO MI benchmark (mean over five seeds; seed s.d. *≤* 1.5 pp), comparing hTSMNet with the strongest comparator, Deformer, as geometric and adaptation components are added left to right. Columns 1–4 are evaluated *without* test-time AdaBN (the canonical main-text footing): the first three are strict zero-shot (*k*=0) and isolate the frontend, and the fourth adds few-shot classifier-only calibration (*k*=20). The final column isolates the test-time AdaBN lever (zero-shot, *k*=0, AdaBN on). “Fixed RBF only” is a geometry-only projection onto the *D*=15 canonical anchors (for Deformer, the fixed comparator projection of Section 4.3; for hTSMNet, the no-correction frontend *A_s_*=polar(*W* ^rbf^)—the two are not identical, see Section 4.3). The learned per-anchor RBF width and the learned MLP correction are specific to the hTSMNet frontend and have no analogue for a fixed-input network (“n/a”). The comparator is thus already given the hTSMNet geometric coordinate system. hTSMNet’s advantage comes from learning the alignment and coupling it to the SPD backbone.

| Model | Zero-shot (*k*=0): frontend build-up |  |  | Adaptation (full model) |  |
|---|---|---|---|---|---|
|  | Fixed RBF only | + learned RBF width | + learned MLP (full frontend) | *k*=20 classifier calibration | + test-time AdaBN (*k*=0) |
| hTSMNet | 56.9 | 64.3 | 67.3 | **75.7** | **71.9** |
| Deformer | **66.4** | n/a | n/a | 72.4 | 70.2 |

## 3 Discussion

### 3.1 Summary

We introduced hTSMNet, a geometry-conditioned hyperalignment frontend that maps any variable-montage recording into a shared anchor space and feeds it to a Riemannian SPD backbone. Because the alignment is generated from sensor coordinates alone, hTSMNet transfers zero-shot to unseen subjects, datasets, and montages. Three findings follow. (i) On a leave-onedataset-out MI benchmark spanning three datasets and 172 subjects, hTSMNet is competitive at zero-shot and attains the best few-shot accuracy on every dataset, with a per-subject advantage that is significant against all eight comparators (Section 2.3). (ii) The reason is geometric: hTSMNet learns an LH/RH class direction shared across datasets, attaining the highest crossdataset class-direction consistency of any method (0.71 cosine, versus 0.21 with no learning), so a single decision direction is already oriented correctly on a held-out dataset (Section 2.4). (iii) A *frozen* MI model transfers across paradigms along its band-power substrate, decoding workload (84%) and SSVEP (68–72%) but not P300, a dissociation confirmed by a frozen-probe and scalp-pattern analysis (Sections 2.6–2.7). We discuss the practical boundaries of these claims below. Together these trace one thread, from a geometry-conditioned shared anchor representation, through cross-dataset transfer and a stable tangent-space representation, to cross-paradigm transfer within the oscillatory regime.

### 3.2 What the alignment learns: a shared coordinate frame, not a leadfield inverse

A natural misreading of hTSMNet, first flagged in Section 2.1, is that the frontend inverts the forward model—that *A_s_*approximates the leadfield inverse 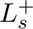 and the anchors are recovered sources. It does neither, by design. The leadfield matrix *L_s_* of Equation (1) is the true physical source-to-sensor projection: subject-specific and unknown. Inverting it, 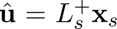, is ill-posed and subject-specific—a far harder problem than the one we need to solve. The frontend’s job is narrower: it only makes the downstream anchor coordinates comparable across recordings. What *A_s_* actually shapes is the composite 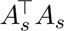 of Equation (2)—the net source-to-anchor map—never *L_s_* itself, which the frontend never sees. Because *A_s_* is generated from one shared function of geometry, the training objective encourages this composite toward a single source-to-anchor operator Π common across recordings, so that an anchor coordinate carries comparable source signal across subjects and montages without ever estimating *L_s_* or its inverse. The anchors are therefore best read as a reusable *surrogate* for a shared source geometry—fixed reference locations that stand in for “where” a coordinate lives—rather than as recovered cortex. This is a deliberate trade: by giving up on source reconstruction we obtain a map that is well posed, montage-agnostic, and parameter-free across subjects. That the composite does converge to a common frame in practice, rather than merely in principle, is what the cross-subject receptivefield consistency (Section 2.2) and the shared class direction (Section 2.4) demonstrate.

### 3.3 Relationship to variable-montage foundation models

hTSMNet is closest in spirit to variable-montage models that use sensor positions to map heterogeneous recordings into a shared space, such as LUNA [9] and LaBraM [10]. The distinction is that hTSMNet makes the alignment *explicit and inspectable*. The frontend emits a linear operator *A_s_* that can be visualised as per-anchor receptive fields (Figure 2D), reflecting the underlying source-to-sensor (leadfield) geometry, and shown to be consistent across montages (Figures 2E–F), rather than leaving alignment implicit in attention weights. The anchors are physical scalp locations with a defined geometry, and the aligned signal is processed on the SPD manifold, preserving the covariance structure that MI decoding depends on. The head-tohead LODO comparison (Section 2.3) indicates that, in the small-to-medium MI regime studied here, this combination of explicit geometric alignment and Riemannian processing outperforms large position-aware transformers. We caution that these transformers were developed for broad self-supervised pretraining rather than this specific small-MI regime, so the comparison reflects fitness for the present task rather than a general ranking.

### 3.4 Why a Riemannian SPD backbone

We deferred from Section 2.1 the question of why the geometry-conditioned frontend is paired with TSMNet rather than a generic convolutional or transformer decoder. There are three reasons. First, *physiological fit*: the discriminative content of motor imagery is carried by oscillatory power and its spatial covariance, exactly the second-order structure that SPD and tangent-space methods are built to exploit [11, 12]. Second, *sensitivity to the frontend*: the learned correction re-weights which sensors feed each anchor, but this only helps if the decoder turns such changes into discriminative features. The cross-backbone transplant of Section 2.5 shows it does so far more on a Riemannian than on a convolutional decoder—learning the alignment yields +10.4 pp on the deep SPD backbone but only +3.3 pp on EEGNet. The reason is that covariance pooling followed by the LogEig map is a strongly nonlinear readout in which small, geometry-consistent changes to the alignment are magnified into large tangentspace differences—so the SPD backbone is not merely compatible with the geometric frontend, it is the component that converts the frontend’s freedom into accuracy. Third, *interpretability*: operating on covariances of a fixed anchor space lets a trained model be traced back to cortex and frequency—each anchor maps to a scalp location and each temporal filter to a band—which is what makes the cross-paradigm scalp-pattern traceback of Section 2.7 possible. Together these motivate pairing the hyperalignment frontend with a Riemannian, rather than a purely convolutional, decoder.

This choice also connects hTSMNet to the broader theory of SPD-domain adaptation. Li *et al.* [14] show that SPD/tangent-space methods offer invariance to linear mixing, a property well suited to EEG, and that source-free adaptation of their second-order statistics can compensate conditional and label distribution shifts on the SPD manifold. This provides the closest theoretical context for the label-free SPD batch-norm adaptation used here.

### 3.5 The cross-paradigm boundary is informative

The cross-paradigm result is central to the study. Its value lies less in the raw transfer numbers than in what they reveal about *generalization*. A network trained only to tell left hand from right—on a small, single-paradigm corpus of three MI datasets—is a generalizable extractor for oscillatory band-power contrasts. Mental workload and SSVEP, paradigms hTSMNet never saw and that bear no behavioural resemblance to motor imagery, are recovered by re-fitting nothing but the linear readout, while ERP/P300 cannot be. The relevant unit of generalization is therefore not the named paradigm but the *signal family* the backbone represents. Read this way, the P300 result constrains the claim rather than contradicting it: the model’s reach is *largely predictable in advance* from a single property of the target signal (band-limited vs. phaselocked), so the scope statement is a positive specification of where the representation applies, not an unexplained average over tasks.

This predictability has a direct practical consequence for calibration. The frozen-probe ceiling (Section 2.7) is not only a retrospective explanation but a *pre-adaptation diagnostic*: faced with a new task, one can ask whether its discriminative contrast is already linearly present in the frozen representation before committing to data collection or fine-tuning. A high probe signals that a short labelled support block should suffice to fit a task-specific classifier. A chance-level probe warns that classifier adaptation alone is unlikely to recover the task signal. For the broad family of oscillatory, passive-BCI applications—workload, fatigue, vigilance, steady-state stimulation, and candidate clinical screening—this turns a single MI-pretrained, montage-agnostic model into a label-efficient starting point rather than a motor-imagery-only classifier, with calibration reduced to a re-fit classifier on a handful of trials and no per-paradigm pretraining.

Finally, the finding bears on how generalizable EEG models are built. Cross-paradigm reuse—at least within the oscillatory regime—need not come only from massive, multi-paradigm self-supervised pretraining: a comparatively small supervised MI corpus already yields it when the representation carries the right inductive bias, montage-normalised covariance on a shared anchor space. We do not read this as an argument against large EEG foundation models, nor as evidence that hTSMNet covers phase-locked paradigms as built. Rather, it identifies what a broadly generalizable decoder must *preserve*—the band-power geometry that already unifies MI, workload, and SSVEP—while motivating a natural extension: a phase-preserving evoked-template branch, run alongside the covariance branch and feeding the same shared-anchor machinery, so the model can also carry the trial-aligned structure ERP tasks depend on.

### 3.6 Limitations and future work

Several caveats bound our conclusions.

- *Paradigm scope of the empirical claims.* The architecture is paradigm-agnostic, and the cross-paradigm experiments extend validation beyond MI for the first time, but the model trained here reads only band-power. ERP, sleep, and clinical paradigms whose signal is phaselocked would require a phase-preserving evoked-template branch feeding the same sharedanchor machinery. A direct future comparison should bracket the frozen-MI plus new-classifier transfer result against models fully trained on the target paradigm. For workload, the withintarget EA–TSLR upper reference already gives one such marker at 91.3%.
- *Rank mismatch. D >* min*_s_ N_s_* makes the aligned covariance rank-deficient. Spherical-spline upsampling to virtual channels or channel-dropout augmentation are principled mitigations we leave to future work. A controlled same-dataset channel-ablation probe, progressively dropping channels from a dense montage while holding the dataset fixed, would separate channel-count effects from cross-dataset shift.
- *Anchor template as a design choice.* We treat anchor positions as a fixed, interpretable, reusable design artefact rather than learning them end-to-end. Joint learning of anchors invites failure modes (collapse onto sensor-dense regions, drift outside cortically meaningful territory, or specialisation to the dominant training dataset), each of which would erode the cross-dataset canonicality that motivates the architecture, for an expected accuracy gain that Figure S2 suggests is small. Coverage-aware template construction on heterogeneous corpora, and a characterisation of when a universal cross-corpus template suffices versus a corpus-specific one, are the more promising directions.
- *Binary sub-task.* All real-data results use the binary LH/RH (or two-class) decision. The backbone already supports more than two classes, so multi-class hTSMNet is a direct extension, but the representational-geometry metrics reported here are defined for the binary case.
- *Comparator coverage.* The deep geometric/SPD family is represented here by the TSMNet backbone and its shallow tangent-space variant (Figure 5). A dedicated external geometricdeep-learning baseline (*e.g.* an SPD network or Tensor-CSPNet) is left for future work.
- *No source localisation.* The anchors are reference scalp locations and the traceback (Section 2.7) yields sensor-space topographies, not cortical source estimates. We make no inverseproblem claim, and the neurophysiological correspondences we report are interpretive.
- *Preprocessing dependence.* All data pass through one MOABB pipeline (fixed bandpass, [0, 2] s window, `standard_1020` channel matching, common-average reference). Robustness to alternative epoching, referencing, and channel-matching conventions, particularly for the ERP and SSVEP targets, remains to be characterised.
- *Statistics.* Finally, our error bars report seed variability and the per-subject tests pool subjects across folds. A hierarchical mixed-effects analysis that separates seed, support-draw, and between-subject variance would tighten the uncertainty statements.

## 4 Methods

### 4.1 hTSMNet architecture

hTSMNet has two modules trained end-to-end: a *hyperalignment frontend* that maps any variable-montage input to a fixed-*D* anchor space, and the *TSMNet backbone* [13] that classifies in that space.

#### Geometry

Electrode coordinates are canonical standard 10–20 positions with the 10–05 extension, looked up via MNE’s `standard_1020` montage from channel labels and not measured per subject. Deployment therefore requires correct channel labels, not a digitiser. Electrode and anchor coordinates are reduced to 2D by an orthographic (top-down *x*–*y*) projection of those 3D canonical positions, centred on their median and rescaled so the median sensor radius is 1. Anchors are normalised in the same frame. Anchors are held fixed throughout training. For the headline configuration we place *D* = 15 anchors by farthest-point sampling of the same canonical `standard_1020` layout (a data-agnostic template). The *k*-means, random, and paradigm-specific (sensorimotor) templates used in Figure S2 are constructed analogously.

#### Alignment matrix

Given sensor positions 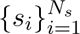 and anchors 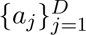, the alignment is

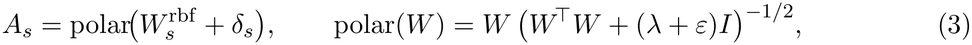

with Tikhonov damping *λ* = 10*^−^*^3^ (the regulariser) and a numerical floor *ε* = 10*^−^*^6^ that keeps the inverse square root well defined even as *λ →* 0. The two enter only through the single positive shift *λ* + *ε*. The damped polar retraction is a single continuous formula valid for both *N_s_ ≥ D* and *N_s_ < D* (as *λ →* 0 with *N_s_ ≥ D* it recovers the standard column-orthonormal polar), avoiding a discrete branch swap that would perturb anchor identity at low rank. The RBF base is 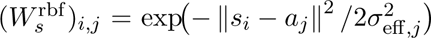 with a learned per-anchor width 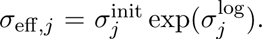 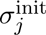 is *data-conditioned*: the median distance from anchor *j* to its nearest sensors over the union of source-domain sensor clouds (clamped to [0.05, 2.0] in normalised units), so each anchor’s receptive scale matches local sensor density. The log-multiplier 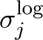 is learned (initialised to 0, so 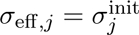 at start) under a soft *L*_2_ prior (*λ_σ_* = 10*^−^*^3^) that keeps it near the data-conditioned reference.

#### Correction generator

The correction *δ_s_∈* ℝ*^N_s_×D^* is produced by a single network shared across all subjects. For each (sensor *i*, anchor *j*) pair it concatenates Fourier position embeddings of *s_i_* and *a_j_* (random Fourier features with 16 learnable frequencies projected to 64 dimensions), the difference *s_i_ − a_j_*, the distance ||*s_i_* − *a_j_*||, and the scalar 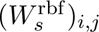, and passes them through a 2-layer MLP (hidden 64, GELU, dropout 0.1) to a scalar *δ_i,j_*. This MLP’s output layer (the *frontend head*, not to be confused with the downstream linear classifier) is initialised with a small Gaussian weight (standard deviation 10*^−^*^3^, zero bias), so at the first step *δ ≈* 0 and *A_s_ ≈* polar(*W* ^rbf^)—the frontend is RBF-dominant at warm-start and grows the learned correction only where the task earns it (setting this output-layer weight to exactly zero gives *δ ≡* 0 and defines the geometry-only RBF ablation of Figure 5). The frontend head is learned only during source pretraining and is held frozen at few-shot calibration time, which re-fits the linear classifier alone. *The frontend has no per-subject parameters*: the only learned objects are the shared generator weights and the per-anchor *σ*^log^, so the alignment of any recording (seen or unseen) is a deterministic function of its sensor coordinates. This is what makes the zero-shot forward path (Section 2.1) possible.

#### TSMNet backbone

The anchor-space signal **z***_s_∈* ℝ*^D×T^* is processed by the TSMNet pipeline [13], which we group into three stages (*f_θ_*, *m_ϕ_*, *g_ψ_*). The feature extractor *f_θ_* applies a depthwise temporal convolution followed by a *D ×* 1 spatial convolution. The SPD-manifold stage *m_ϕ_* pools the channel covariance into an SPD matrix, reduces and rectifies it with a BiMap bilinear layer (which lowers the SPD dimension from the number of spatial filters to a smaller *d*_out_ *× d*_out_ subspace, with *d*_out_ = 32 on all three MI LODO folds) and a ReEig eigenvalue floor, applies a shared SPD batch-normalisation that whitens by the running Riemannian (Fréchet) mean of the mini-batch covariances, and maps the result to the tangent space at the identity with a LogEig layer, whose vectorised upper triangle—of dimension *d*_out_(*d*_out_+1)*/*2 (= 528 for all three folds)—is the feature the classifier sees. The classifier *g_ψ_*is a single linear layer on the resulting tangent vector. We use the *shared* (not domain-specific) SPD batch-norm, which is compatible with zero-shot inference for unseen subjects. A domain here is a single subject (we did not subdivide by session). All SPD operations and their custom backward passes run in double precision for eigendecomposition stability. In all runs reported here, training additionally includes a self-supervised covariance-alignment term—the mean log-Euclidean distance between the per-domain anchor-space time-covariances over the domain pairs present in each multi-subject mini-batch—weighted at *λ*_cov_ = 0.05 (covariance floor 10*^−^*^6^, shrinkage 0.01). It is a *class-agnostic* auxiliary regulariser that nudges each domain’s anchor-space second-order statistics toward a common distribution. Setting *λ*_cov_ = 0 removes it.

#### Training

The pooled source data are split 80*/*20 into training and validation sets, and all parameters (frontend and backbone alike) are trained jointly with a single Adam optimiser (lr = 3 *×* 10*^−^*^4^, weight decay 10*^−^*^3^, batch size 256, gradient clipping at norm 5) under a cross-entropy loss—augmented by the auxiliary covariance-alignment term and the soft *σ*^log^ prior noted above— with early stopping when source-validation accuracy stops improving. The shared generator is given a 5*×* larger learning rate than the rest of the network (a single optimiser with two parameter groups). We deliberately use a *standard* (non-Riemannian) optimiser throughout: the manifold constraint on *A_s_* is enforced at forward time by the polar retraction of Equation (3), and the SPD layers keep their iterates on the manifold by construction (the ReEig eigenvalue floor), so no Riemannian optimiser is needed—which also avoids the eigendecomposition instabilities that Riemannian updates on SPD parameters can introduce. Inputs are normalised per trial by their global standard deviation. For an unseen subject the alignment is computed from sensor positions alone. *Test-time AdaBN is off by default*: in the main LODO benchmark (Section 2.3) and in the cross-paradigm transfer (Section 4.5) we report results with no targetdomain adaptation, so every method is under the same evaluation settings and *k* = 0 is strict zero-shot—no target labels and no target statistics. The optional AdaBN step—recomputing the shared SPD batch-norm statistics from a held-out subject’s *unlabelled* trials (an unsupervised Fréchet-mean update)—is a label-free, transductive variant studied separately in Section 2.8 and Figure S3. It is the only place AdaBN is enabled. hTSMNet’s SPD batch-norm recomputes the running mean as the Riemannian (Fréchet/Karcher) mean of the anchor-space covariances on the SPD manifold, whereas the convolutional BatchNorm comparators (EEGNet, Conformer, Deformer) use the ordinary Euclidean BatchNorm recompute, so AdaBN is matched to each architecture’s normalisation.

All experiments use the lean implementation stack described by the release code: PyTorch *≥* 2.7 with CUDA 12.8 for GPU runs, plus MNE and MOABB for dataset access, preprocessing, and canonical montage lookup.

#### Few-shot adaptation

Few-shot adaptation models the realistic deployment setting in which a new user supplies only a short labelled calibration block before use. Given *k ∈ {*0, 5, 10, 20, 50*}* class-balanced support trials from the target, we adapt only the linear final-layer classifier head (30 Adam steps). The feature extractor is frozen. The model carries two sets of learned weights downstream of the input—the correction MLP *inside* the geometry frontend (the “frontend head”, defined above) and the final tangent-space linear classifier (the “final-layer classifier”)— and few-shot calibration touches *only the latter*: the frontend, including its MLP, and the SPD backbone stay frozen, so adaptation never changes the montage alignment, only the read-out. *k* = 0 is the strict zero-shot forward pass. Our headline operating point is the few-shot regime (*k* = 20, a short calibration block), where hTSMNet’s advantage is clear. The upper end (*k* = 50 per class) is deliberately generous—larger than a typical EEG calibration block—and is included to show that the method ranking is stable as the curves saturate, not as a recommended budget. Five independent support draws are averaged per (*k,* subject). The same classifier-only protocol—30 Adam steps adapting only the linear classifier with the body frozen—is applied to every method. Following common practice, each method’s classifier learning rate is set to its own tuned value (hTSMNet 10*^−^*^2^ with an L2-SP anchor to the source classifier; comparators in the 10*^−^*^4^–10*^−^*^3^ range), so that every method is adapted under a favourable setting rather than a single shared step size.

### 4.2 Datasets and preprocessing

All data are accessed through MOABB [20] and harmonised into one on-disk corpus (Table 2). For each subject we query the relevant MOABB paradigm with a 0.5–45 Hz zero-phase Butterworth bandpass, resampling to *f_s_* = 250 Hz, and a [0, 2] s post-cue window (*T* = 500 samples). Electrode labels are mapped to `standard_1020` 3D positions and unresolved channels are dropped. A minimum of 8 matched electrodes per subject is applied uniformly across the corpus; every pooled MI training subject retains at least 13 matched channels, so this threshold removes no MI subject while preserving the low-channel cross-paradigm targets. The cross-paradigm transfer *targets* are evaluated on their native montages regardless of channel count—including the 8-channel Kalunga2016 and BNCI2014-008, which thereby fall in the low-rank regime discussed in Section 2.8. The MI corpus comprises 172 subjects after `standard_1020` matching (9*/*9 BNCI2014-001, 54*/*54 Lee2019-MI, 109*/*109 PhysionetMotorImagery; none excluded). Two smaller counts reported elsewhere are subsets of this total. Each LODO fold splits its pooled source 80*/*20 at the subject level into training and validation. For the held-out-BNCI2014-001 fold this gives 130 source-training subjects (43 Lee2019-MI + 87 PhysionetMotorImagery) and 33 validation subjects. The frontend receptive-field analysis (Figures 2E–F) is montage-level rather than subject-level: *A_s_* depends only on a montage’s sensor geometry, so the fold has just three distinct alignments—the two source montages and the zero-shot BNCI2014-001 target. The persubject paired test (Figure 3D–E) pools 171 of the 172 subjects: one PhysionetMotorImagery subject has too few trials in a class to leave a held-out query set after the *k*=20 (20-per-class) support draw, so it has no *k*=20 score—the same trial-budget limit that removes Physionet’s *k*=50 operating point. A common-average reference is applied, and the retained 3D positions are stored with each trial so that the frontend (and the position-aware LUNA baseline) consume consistent geometry. We deliberately apply the *same* bandpass and [0, 2] s window to every paradigm: the cross-paradigm dissociation (Section 2.6) then reflects what the frozen representation carries rather than paradigm-specific epoching choices. Crucially, each subject keeps its native montage—no re-referencing to a fixed template.

**Table 2:** Datasets. The three MI datasets form the LODO benchmark and the pooled pretraining corpus. The lower block lists the cross-paradigm transfer targets. Channel counts are after `standard_1020` matching. All data are resampled to 250 Hz.

| Dataset | Paradigm | Subjects | Channels | Two-class sub-task |
|---|---|---|---|---|
| BNCI2014-001 | MI | 9 | 22 | left vs. right hand |
| Lee2019-MI | MI | 54 | 56 | left vs. right hand |
| PhysionetMotorImagery | MI | 109 | 64 | left vs. right hand |
| Hinss2021 | Workload | 15 | 61 | low vs. high workload |
| Lee2019-SSVEP | SSVEP | 54 | 56 | stimulus-frequency contrast |
| Kalunga2016 | SSVEP | 12 | 8 | stimulus-frequency contrast |
| BrainInvaders2013a | P300/ERP | 24 | 16 | target vs. non-target |
| BNCI2014-008 | P300/ERP | 8 | 8 | target vs. non-target |

The cross-dataset LODO benchmark uses three MI datasets (BNCI2014-001 [27], Lee2019MI [28], PhysionetMotorImagery [29, 30]; 172 subjects pooled), restricted to the binary LH/RH sub-task. The cross-paradigm experiments add five datasets from three unseen paradigms— workload (Hinss2021 [31]), SSVEP (Lee2019-SSVEP [28], Kalunga2016 [32]), and P300/ERP (BrainInvaders2013a [33], BNCI2014-008 [34])—each evaluated as a balanced two-class sub-task with a freshly fitted binary classifier.

### 4.3 Comparator methods

We benchmark against eight comparators in three families, each reproduced with its authorrecommended training recipe and adapted to the protocol only where strictly necessary (source training over the pooled datasets; the classical and specialist comparators are projected onto the same *D* = 15 canonical anchors by a fixed RBF interpolation, detailed below under *Comparator input projection*, while the foundation models keep their own fixed-channel or native montages; see Table 3). *Classical Riemannian pipelines*: CSP–LDA (one-vs-rest common spatial patterns with shrinkage LDA [21]) and EA–TSLR (Euclidean alignment [3] followed by tangent-space logistic regression [11, 12]). Both re-fit on the target support set. *Specialist deep networks*: EEGNet [22], EEG-Conformer [23], and EEG-Deformer [24], pretrained on the projected source data. *Pretrained foundation models*: MIRepNet [25] and NeuroGPT [26] (loaded from their official checkpoints—the publicly released pretrained weights—and, for the LODO MI benchmark, fine-tuned on the source MI data through their fixed-channel templates), and LUNA [9], a variable-montage position-aware transformer evaluated on each subject’s native montage. (In the cross-paradigm experiments the foundation models are instead kept at their official pretrained checkpoints; see Section 4.5.) At test time all methods share the classifier-only *k*-shot protocol of Section 4.1. For the classical pipelines, adaptation reduces to re-centring the target covariances and re-fitting the linear decoder, matching their intended use. Input handling therefore differs by comparator family, and we summarise it explicitly in Table 3.

**Table 3:** Input handling per comparator (LODO MI benchmark). “RBF(*D*=15)” is the fixed RBF interpolation onto the *D* = 15 canonical anchors defined under *Comparator input projection* in Section 4.3 (a single global width, no polar retraction; not identical to hTSMNet’s geometryonly ablation). The label “native montage” means the method consumes each subject’s own channels and positions, and “own template” means the model’s published fixed-channel template. In the cross-paradigm experiments the foundation models are kept at their official pretrained checkpoints (Section 4.5). Classical and specialist comparators with RBF(*D*=15) are therefore already given the hTSMNet geometric coordinate system.

| Method | Family | LODO MI input | Source pretraining |
|---|---|---|---|
| hTSMNet | ours | native (geometry-conditioned) | pooled source MI |
| CSP–LDA | classical | RBF(*D*=15) | none (re-fit on target) |
| EA–TSLR | classical | RBF(*D*=15) | none (re-fit on target) |
| EEGNet | specialist | RBF(*D*=15) | pooled source MI |
| EEG-Conformer | specialist | RBF(*D*=15) | pooled source MI |
| EEG-Deformer | specialist | RBF(*D*=15) | pooled source MI |
| NeuroGPT | foundation | own template | official ckpt + source fine-tune |
| MIRepNet | foundation | own template | official ckpt + source fine-tune |
| LUNA | foundation | native montage | official ckpt + source fine-tune |

#### Comparator input projection

The classical and specialist comparators cannot ingest a variable montage, so each native recording is first mapped onto the shared anchor space by a *fixed, unlearned* radial-basis interpolation. We build a static projector *P ∈* ℝ*^D×Ns^* with entries 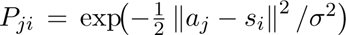 for anchor *a_j_* and sensor *s_i_*, using a *single global* width *σ* = 0.2 in normalised electrode coordinates and *no* row normalisation (an anchor whose total weight underflows falls back to its nearest sensor). The comparator then sees the projected trial *P* **x***_s_ ∈* ℝ*^D×T^*. The anchors 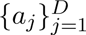 are the same *D* = 15 farthest-point `standard_1020` template hTSMNet uses. This is deliberately the *simplest* montage bridge that still places every comparator in the hTSMNet coordinate system: it is the raw RBF *base* of our frontend and applies *neither* the per-anchor data-conditioned widths *nor* the polar (near-orthonormal) retraction of Equation (3). It is therefore *not* identical to hTSMNet’s geometry-only ablation *A_s_* = polar(*W* ^rbf^) (the no-correction variant in Figure 5), which adds both. Giving the fixedinput comparators this geometric projection is intentionally generous—it lets them decode across heterogeneous montages at all, so the remaining gap to hTSMNet reflects the learned alignment and the SPD backbone rather than the montage bridge itself.

### 4.4 Representational-geometry metrics

The analyses in Section 2.4 use *per-dataset class centroids* of each method’s penultimate features— for hTSMNet the LogEig tangent-space vector (528-dimensional on all three MI folds), for the comparators their pre-classifier features. Features are first *z*-scored per dimension across the pooled trials of the fold (so the metric is scale-free across methods of different latent dimension). For each dataset *d* and class *c ∈ {*LH, RH*}* the centroid 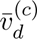 is then the mean of the standardised features over that dataset’s class-*c* trials. The metric is computed per LODO fold (held-out dataset vs. the two sources) and averaged over the three folds.

The *class-direction consistency* measures how similarly the LH/RH discriminative direction is oriented across datasets. For each dataset *d* we form the unit class axis 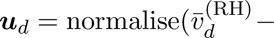 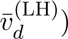 from the *z*-scored centroids and define

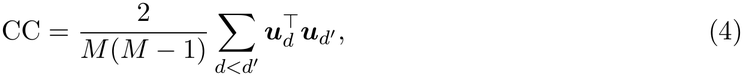

the mean pairwise cosine over the *M* datasets of a fold (here *M* = 3: the held-out dataset and the two sources; *M* is the dataset count, distinct from the anchor dimension *D*). CC *→* 1 means the class axis points the same way in every dataset, so a single linear rule transfers. CC *→* 0 means the class directions are unrelated. Because CC depends only on the *angle* between class axes, it is invariant to feature scale and latent dimension and cannot be improved by rescaling or compressing the representation. In Figure 4A,B the horizontal axis is the frozen *k* = 0 class direction and the vertical axis is the dataset-centroid direction, so each panel shows the shared class axis (whose cross-dataset agreement CC summarises) against the residual dataset offset. We note that the pretrained foundation models are omitted from this metric because their penultimate representations are not produced in a shared, fixed-dimensional space, so the per-dataset class-axis cosine is not comparable across them.

### 4.5 Cross-paradigm transfer and frozen-representation analyses

#### Transfer protocol

Each method is trained once on the pooled three-dataset MI corpus (LH/RH), persisted, and transferred to every cross-paradigm target by freezing the entire network and fitting only a fresh binary classifier on *k* support trials per class (five seeds; AdaBN disabled, so the test is purely inductive). The classifier is fitted separately for each target subject: each seed draws *k* support trials per class from that subject, trains a fresh classifier, and scores the subject’s remaining trials. There is no session split or pooled multi-subject target classifier. For the foundation models the frozen body is their official pretrained checkpoint (not the MI-fine-tuned LODO variant), since for them the transferable representation is the foundation pretraining itself. As for every other method, only a fresh classifier is fitted. EA–TSLR re-fits its full alignment and decoder on the target and is therefore reported as a within-target upper reference rather than a transfer method (omitted from Figure 6).

#### Frozen probe (“signal present?”)

For Figure 7A we train a shrinkage-LDA probe (5-fold within-subject cross-validation) on the frozen MI latent of each target and report balanced accuracy. This measures whether the target’s discriminative signal is linearly present in the frozen representation, independent of any classifier adaptation.

#### Band-power vs. evoked decoding

For Figure 7B we decode the class label from two feature views of the same trials: band-power/covariance features (the representation hTSMNet’s pooling sees) and evoked/time-domain features (the trial-averaged, phase-locked waveform), each scored by balanced accuracy. The contrast isolates whether a paradigm’s signal is oscillatory (recoverable by covariance) or phase-locked (discarded by covariance).

#### Scalp-pattern traceback

For Figure 7D–G we fit a CSP on the frozen latent of each paradigm, take the more spatially focal of the two dominant components (*u*), and back-project it through the frozen frontend onto the scalp to obtain a sensor-space pattern (top-3 sensors annotated). The spectral stem below each topography places every learned temporal filter at its peak frequency (horizontal axis). The stem height *|α_ft_|* is that filter’s loading on the dominant source, the quadratic form 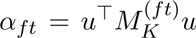 of *u* restricted to temporal filter *ft*, where 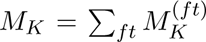 is the anchor-space discriminative matrix traced back from the decoder (reported on an unnormalised relative scale). This identifies, on the scalp and in frequency, the pattern the frozen model reads for each paradigm. These are sensor-space topographies, not cortical source estimates. We do not perform source localisation.

## Data and code availability

All datasets are publicly available through MOABB (https://github.com/NeuroTechX/moabb). Release of the hTSMNet implementation is subject to institutional and intellectual-property review. Details will be provided to the extent permitted upon publication.

## Acknowledgements

This paper is based on results obtained from a project, JPNP14004, commissioned by the New Energy and Industrial Technology Development Organization (NEDO). H.H was supported by Grants-in-Aid for Transformative Research Areas (22H05160) and Raynor Cerebellum Project. O.Y was supported by Japan Agency for Medical Research and Development AMED (JP24wm0625401s0202, JP23wm0625001).

## Supplementary Figures

**Figure S1:**
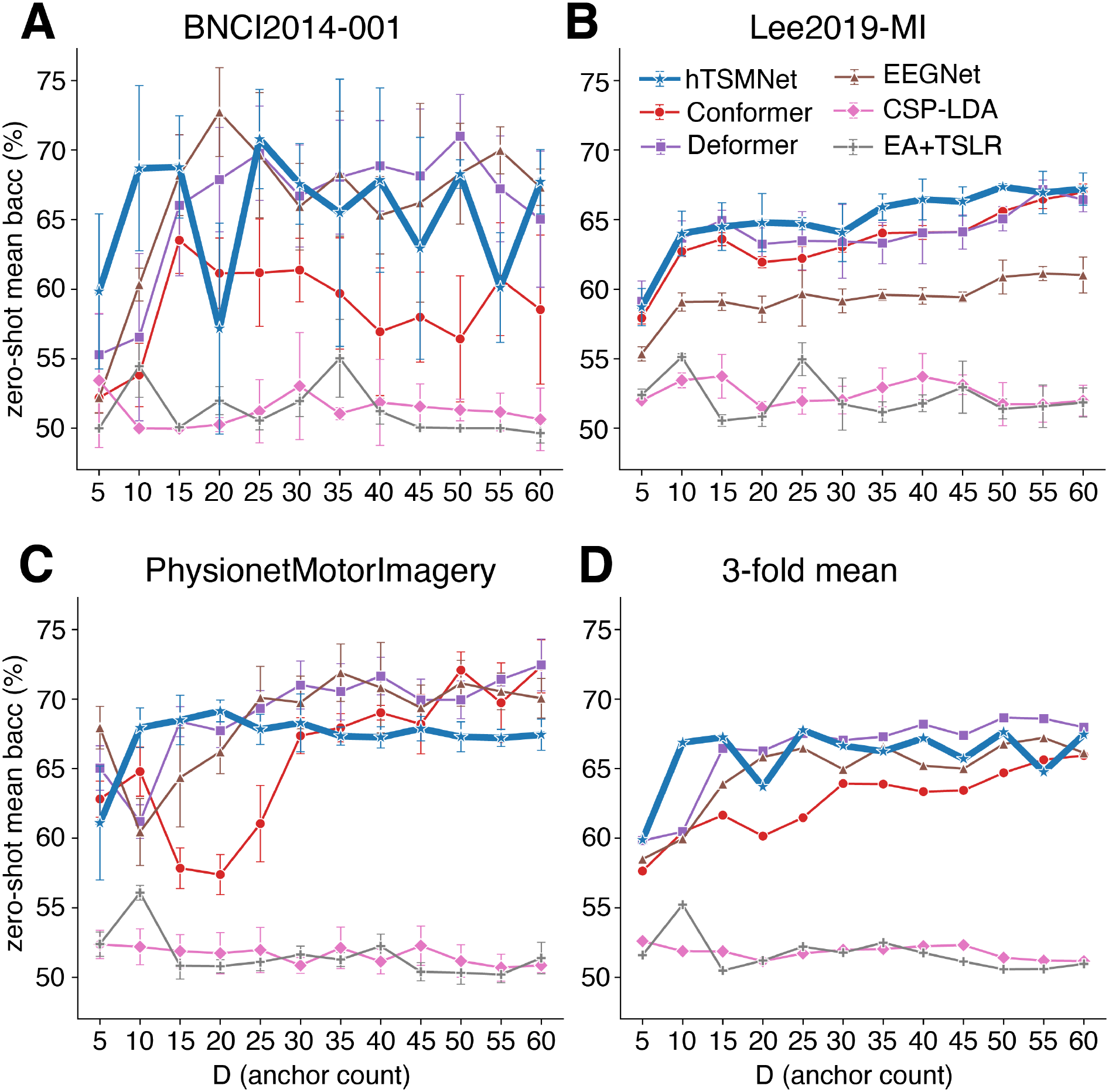
Zero-shot LODO accuracy versus anchor count. *D*. Each panel shows one held-out dataset and the three-fold mean as the anchor dimension is swept over *D ∈ {*5*, …,* 60*}*. Accuracy rises steeply to *D ≈* 15 and then saturates for every method, which motivates the *D*=15 default used throughout. At strict zero-shot hTSMNet (blue) is competitive with the strongest specialist deep networks across the sweep—leading on BNCI2014-001 but not uniformly on Lee2019-MI or PhysionetMotorImagery (cf. Section 2.3).

**Figure S2:**
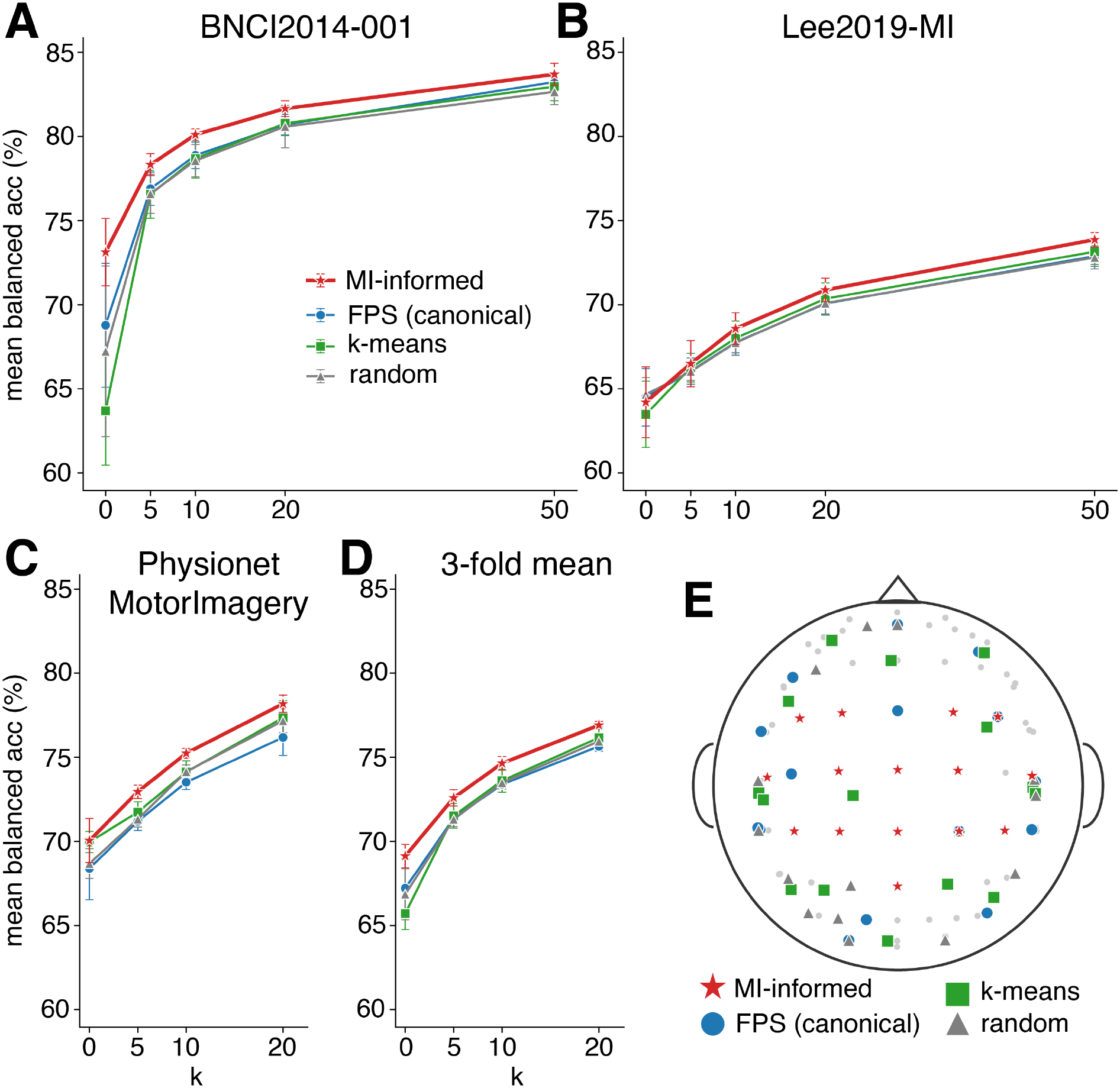
**Robustness to the anchor template (***D* = 15**). (A–D)** Few-shot LODO accuracy for four anchor-template constructions on each dataset and the three-fold mean. The dataagnostic templates—the canonical farthest-point sampling default, *k*-means of the pooled sensor cloud, and a random draw—fall within *∼* 1.5 pp of one another at every *k*. A paradigm-informed (sensorimotor) template adds only *∼* 1.9 pp at zero-shot on the three-fold mean. **(E)** The four anchor templates on the scalp. hTSMNet’s accuracy is thus insensitive to exact anchor placement (Section 2.8).

**Figure S3:**
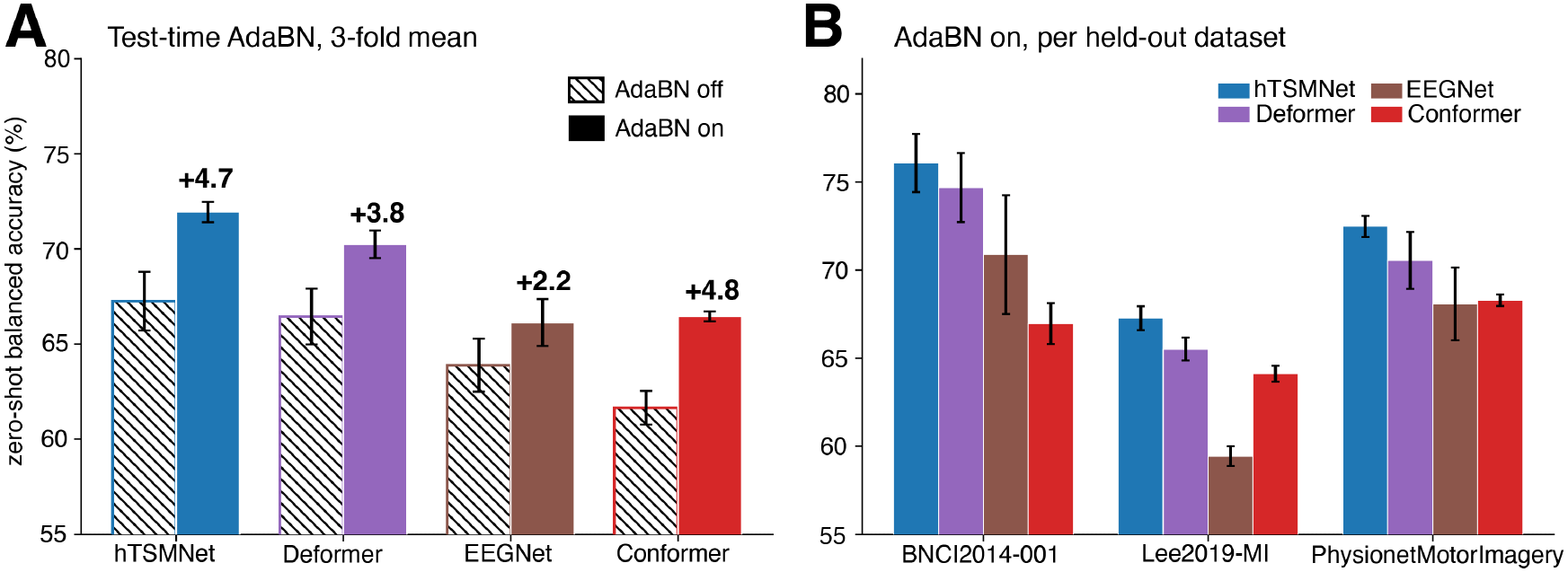
Test-time AdaBN and the zero-shot LODO ceiling for BatchNorm decoders (Section 2.8). AdaBN recomputes a network’s BatchNorm running statistics from each held-out subject’s *unlabelled* trials—a test-time step on the same frozen checkpoints, with no labels and no weight updates—applied uniformly to the four BatchNorm decoders. **(A)** Zeroshot (*k*=0) three-fold-mean balanced accuracy, AdaBN off (hatched) vs. on (solid); mean*±*std over five seeds, gain annotated. AdaBN lifts every decoder (+2.2 to +4.8 pp); hTSMNet stays highest at 71.9%, ahead of Deformer 70.2%. **(B)** The same AdaBN-on accuracies per held-out dataset; hTSMNet (blue) leads on all three. Non-BatchNorm baselines (CSP–LDA, EA–TSLR, LUNA) cannot use AdaBN and are omitted.

## References

[1] F. Lotte, L. Bougrain, A. Cichocki, M. Clerc, M. Congedo, A. Rakotomamonjy, and F. Yger, “A review of classification algorithms for EEG-based brain–computer interfaces: a 10 year update,” Journal of Neural Engineering, vol. 15, no. 3, p. 031005, 2018.

[2] D. Kostas and F. Rudzicz, “Thinker invariance: enabling deep neural networks for BCI across more people,” Journal of Neural Engineering, vol. 17, no. 5, p. 056008, 2020.

[3] H. He and D. Wu, “Transfer learning for brain–computer interfaces: a Euclidean space data alignment approach,” IEEE Transactions on Biomedical Engineering, vol. 67, no. 2, pp. 399–410, 2020.

[4] P. L. C. Rodrigues, C. Jutten, and M. Congedo, “Riemannian Procrustes analysis: transfer learning for brain–computer interfaces,” IEEE Transactions on Biomedical Engineering, vol. 66, no. 8, pp. 2390–2401, 2019.

[5] T. Ding, Y. Zhang, X. Zhao, J. Shi, J. Liu, Z. Tang, B. Tao, and H. Ding, “RSSDA: a Riemannian source selection and domain alignment framework for cross-subject multi-class motor imagery classification,” Pattern Recognition, 2026, doi:10.1016/j.patcog.2026.113934.

[6] J. Cheng, R. Ran, and B. Fang, “SPD-DANN: an SPD manifold unsupervised domain adaptation method for cross-subject motor imagery EEG decoding,” Neural Networks, 2026, doi:10.1016/j.neunet.2026.109076.

[7] D. Kostas, S. Aroca-Ouellette, and F. Rudzicz, “BENDR: using transformers and a contrastive self-supervised learning task to learn from massive amounts of EEG data,” Frontiers in Human Neuroscience, vol. 15, p. 653659, 2021.

[8] C. Yang, M. B. Westover, and J. Sun, “BIOT: biosignal transformer for cross-data learning in the wild,” in Advances in Neural Information Processing Systems (NeurIPS), 2023.

[9] B. Döner, T. M. Ingolfsson, L. Benini, and Y. Li, “LUNA: efficient and topology-agnostic foundation model for EEG signal analysis,” in Advances in Neural Information Processing Systems (NeurIPS), 2025. arXiv:2510.22257.

[10] W.-B. Jiang, L.-M. Zhao, and B.-L. Lu, “Large brain model for learning generic representations with tremendous EEG data in BCI,” in International Conference on Learning Representations, 2024.

[11] A. Barachant, S. Bonnet, M. Congedo, and C. Jutten, “Multiclass brain–computer interface classification by Riemannian geometry,” IEEE Transactions on Biomedical Engineering, vol. 59, no. 4, pp. 920–928, 2012.

[12] M. Congedo, A. Barachant, and R. Bhatia, “Riemannian geometry for EEG-based brain– computer interfaces: a primer and a review,” Brain–Computer Interfaces, vol. 4, no. 3, pp. 155–174, 2017.

[13] R. J. Kobler, J.-I. Hirayama, Q. Zhao, and M. Kawanabe, “SPD domain-specific batch normalization to crack interpretable unsupervised domain adaptation in EEG,” in Advances in Neural Information Processing Systems (NeurIPS), vol. 35, pp. 6219–6235, 2022.

[14] S. Li, M. Kawanabe, and R. J. Kobler, “SPDIM: Source-free unsupervised conditional and label shift adaptation in EEG,” in International Conference on Learning Representations (ICLR), 2025. OpenReview: https://openreview.net/forum?id=CoQw1dXtGb.

[15] J. V. Haxby, J. S. Guntupalli, A. C. Connolly, Y. O. Halchenko, B. R. Conroy, M. I. Gobbini, M. Hanke, and P. J. Ramadge, “A common, high-dimensional model of the representational space in human ventral temporal cortex,” Neuron, vol. 72, no. 2, pp. 404–416, 2011.

[16] G. Pfurtscheller and F. H. Lopes da Silva, “Event-related EEG/MEG synchronization and desynchronization: basic principles,” Clinical Neurophysiology, vol. 110, no. 11, pp. 1842– 1857, 1999.

[17] A. M. Norcia, L. G. Appelbaum, J. M. Ales, B. R. Cottereau, and B. Rossion, “The steadystate visual evoked potential in vision research: a review,” Journal of Vision, vol. 15, no. 6, p. 4, 2015.

[18] A. Gevins and M. E. Smith, “Neurophysiological measures of cognitive workload during human–computer interaction,” Theoretical Issues in Ergonomics Science, vol. 4, no. 1–2, pp. 113–131, 2003.

[19] J. Polich, “Updating P300: an integrative theory of P3a and P3b,” Clinical Neurophysiology, vol. 118, no. 10, pp. 2128–2148, 2007.

[20] V. Jayaram and A. Barachant, “MOABB: trustworthy algorithm benchmarking for BCIs,” Journal of Neural Engineering, vol. 15, no. 6, p. 066011, 2018.

[21] B. Blankertz, R. Tomioka, S. Lemm, M. Kawanabe, and K.-R. Müller, “Optimizing spatial filters for robust EEG single-trial analysis,” IEEE Signal Processing Magazine, vol. 25, no. 1, pp. 41–56, 2008.

[22] V. J. Lawhern, A. J. Solon, N. R. Waytowich, S. M. Gordon, C. P. Hung, and B. J. Lance, “EEGNet: a compact convolutional neural network for EEG-based brain–computer interfaces,” Journal of Neural Engineering, vol. 15, no. 5, p. 056013, 2018.

[23] Y. Song, Q. Zheng, B. Liu, and X. Gao, “EEG Conformer: convolutional transformer for EEG decoding and visualization,” IEEE Transactions on Neural Systems and Rehabilitation Engineering, vol. 31, pp. 710–719, 2023.

[24] Y. Ding, Y. Li, H. Sun, R. Liu, C. Tong, C. Liu, X. Zhou, and C. Guan, “EEG-Deformer: a dense convolutional transformer for brain–computer interfaces,” IEEE Journal of Biomedical and Health Informatics, vol. 29, no. 3, pp. 1909–1918, 2025.

[25] D. Liu, Z. Chen, J. Luo, S. Lian, and D. Wu, “MIRepNet: a pipeline and foundation model for EEG-based motor imagery classification,” arXiv preprint arXiv:2507.20254, 2025.

[26] W. Cui, W. Jeong, P. Thólke, T. Medani, K. Jerbi, A. A. Joshi, and R. M. Leahy, “NeuroGPT: towards a foundation model for EEG,” in IEEE International Symposium on Biomedical Imaging (ISBI), 2024.

[27] M. Tangermann, K.-R. Müller, A. Aertsen, N. Birbaumer, C. Braun, C. Brunner, et al., “Review of the BCI Competition IV,” Frontiers in Neuroscience, vol. 6, p. 55, 2012.

[28] M.-H. Lee, O.-Y. Kwon, Y.-J. Kim, H.-K. Kim, Y.-E. Lee, J. Williamson, S. Fazli, and S.-W. Lee, “EEG dataset and OpenBMI toolbox for three BCI paradigms: an investigation into BCI illiteracy,” GigaScience, vol. 8, no. 5, p. giz002, 2019.

[29] G. Schalk, D. J. McFarland, T. Hinterberger, N. Birbaumer, and J. R. Wolpaw, “BCI2000: a general-purpose brain–computer interface (BCI) system,” IEEE Transactions on Biomedical Engineering, vol. 51, no. 6, pp. 1034–1043, 2004.

[30] A. L. Goldberger, L. A. N. Amaral, L. Glass, J. M. Hausdorff, P. C. Ivanov, R. G. Mark, et al., “PhysioBank, PhysioToolkit, and PhysioNet: components of a new research resource for complex physiologic signals,” Circulation, vol. 101, no. 23, pp. e215–e220, 2000.

[31] M. F. Hinss, B. Somon, F. Dehais, and R. N. Roy, “Open EEG datasets for passive brain–computer interface applications: lacks and perspectives,” in 10th International IEEE/EMBS Conference on Neural Engineering (NER), pp. 686–689, 2021.

[32] E. K. Kalunga, S. Chevallier, Q. Barthélemy, K. Djouani, E. Monacelli, and Y. Hamam, “Online SSVEP-based BCI using Riemannian geometry,” Neurocomputing, vol. 191, pp. 55–68, 2016.

[33] E. Vaineau, A. Barachant, A. Andreev, P. L. C. Rodrigues, G. Cattan, and M. Congedo, “Brain Invaders adaptive versus non-adaptive P300 brain–computer interface dataset (bi2013a),” Research report, GIPSA-lab, arXiv:1904.09111, 2019.

[34] A. Riccio, L. Simione, F. Schettini, A. Pizzimenti, M. Inghilleri, M. Olivetti Belardinelli, D. Mattia, and F. Cincotti, “Attention and P300-based BCI performance in people with amyotrophic lateral sclerosis,” Frontiers in Human Neuroscience, vol. 7, p. 732, 2013.

